# TALE and NF-Y co-occupancy marks enhancers of developmental control genes during zygotic genome activation in zebrafish

**DOI:** 10.1101/720102

**Authors:** William Stanney, Franck Ladam, Ian J. Donaldson, Teagan J. Parsons, René Maehr, Nicoletta Bobola, Charles G Sagerström

## Abstract

Animal embryogenesis is initiated by maternal factors, but zygotic genome activation (ZGA) shifts control to the embryo at early blastula stages. ZGA is thought to be mediated by specialized maternally deposited transcription factors (TFs), but here we demonstrate that NF-Y and TALE – TFs with known later roles in embryogenesis – co-occupy unique genomic elements at zebrafish ZGA. We show that these elements are selectively associated with early-expressed genes involved in transcriptional regulation and possess enhancer activity in vivo. In contrast, we find that elements individually occupied by either NF-Y or TALE are associated with genes acting later in development – such that NF-Y controls a cilia gene expression program while TALE TFs control expression of *hox* genes. We conclude that NF-Y and TALE have a shared role at ZGA, but separate roles later during development, demonstrating that combinations of known TFs can regulate subsets of key developmental genes at vertebrate ZGA.

## INTRODUCTION

For animal embryogenesis, initial development of the zygote is controlled by parental material deposited into sperm and oocytes during gametogenesis. The bulk of this material is provided by the oocyte and the duration of the maternally controlled period varies between different animal species. However, embryonic development in all animals eventually switches to zygotic control. This maternal-to-zygotic transition (MZT) is a complex process that involves changes in the cell cycle, chromatin state and gene expression (reviewed in (Vastenhouw et al., 2019)). Zygotic genome activation (ZGA) is a key component of the MZT that establishes gene expression programs driving subsequent embryonic development during gastrulation and organogenesis stages. ZGA is thought to be initiated by the action of maternally provided transcription factors (TFs). Accordingly, the Zelda TF induces expression of many genes at ZGA in *Drosophila* (Harrison et al., 2011; Liang et al., 2008; Nien et al., 2011), but there is no known Zelda ortholog in vertebrates. Instead, other TFs have been proposed to regulate ZGA in vertebrates. For instance, Nanog, SoxB1 and Oct4/Pou5f1 (a.k.a. Pou5f3 in zebrafish) are maternally deposited in zebrafish and are required for gene expression at zebrafish ZGA (Lee et al., 2013; Leichsenring et al., 2013). Similarly, Dppa2 and Dppa4 are maternally transmitted and act via Dux TFs to activate large numbers of genes at ZGA in mouse and human embryos (De Iaco et al., 2019; De Iaco et al., 2017; Eckersley-Maslin et al., 2019; Hendrickson et al., 2017). However, subsequent genetic analyses indicate that *Dux* is not required for zygotic development in vivo (Chen and Zhang, 2019) and that the requirement for zebrafish Nanog is likely indirect via an effect on extraembryonic tissues (Gagnon et al., 2018), suggesting that vertebrate ZGA is more complicated and that additional TFs are likely involved in this process.

De novo activation of the zygotic gene expression program requires that the maternally transmitted TFs are able to access their binding sites in compacted genomic DNA, a property associated with pioneer factors (reviewed in (Iwafuchi-Doi and Zaret, 2016)). Accordingly, Zelda opens inaccessible genomic regions to permit binding by other TFs (Schulz et al., 2015; Sun et al., 2015) and both Oct4/Pou5f1 and Sox proteins also possess pioneer activity (Soufi et al., 2012; Soufi et al., 2015). Notably, pioneer factors are active at many stages of embryogenesis to establish tissue-specific gene expression programs. For instance, FoxA TFs control the initiation of hepatic gene expression (Cirillo et al., 2002; Gualdi et al., 1996) and PU.1 controls myeloid and lymphoid development (Heinz et al., 2010). Since the initiation of tissue specific gene expression programs is conceptually similar to zygotic gene activation, it is possible that pioneer factors with known later roles (e.g. in gastrulation or organogenesis) also act at the ZGA. Accordingly, we recently reported that two TFs of the TALE family (Prep1 and Pbx4) – that were originally defined as cooperating with Hox proteins in the activation of tissue-specific gene expression during organogenesis – actually occupy their genomic binding sites already during maternal stages of zebrafish embryogenesis (Ladam et al., 2018). Many of these sites are also occupied by nucleosomes, consistent with previous reports that Pbx TFs possess pioneer activity (Magnani et al., 2011). While we find maternally deposited TALE TFs bound at genomic elements near genes activated at ZGA, they are also bound near genes active later in development (Ladam et al., 2018), suggesting that TALE TFs may have a dual role. Notably, we further demonstrated that a subset of TALE-occupied sites is associated with nearby binding motifs for the NF-Y TF. NF-Y was originally discovered as a ubiquitous basal transcription factor (reviewed in (Dolfini et al., 2012)), but has recently been shown to be maternally deposited in zebrafish, to form protein complexes with TALE TFs and to possess pioneer activity (Ladam et al., 2018; Nardini et al., 2013; Oldfield et al., 2014). Recent work identified NF-Y binding sites near genes activated at ZGA in mouse embryos (Gao et al., 2018; Lu et al., 2016), but NF-Y also binds near many genes acting at later stages of development (Oldfield et al., 2014). Additionally, NF-Y disruption leads to embryonic lethality in mouse (Bhattacharya et al., 2003), but has only mild effects on gene expression at ZGA (Lu et al., 2016).

Here we examine the function of TALE and NF-Y TFs during zebrafish embryogenesis. Our results suggest that these TFs have a shared role at ZGA, but separate roles later during development. In particular, genomic elements individually bound by either NF-Y or TALE TFs are associated with genes acting later in development – such that NF-Y controls a cilia gene expression program while TALE TFs control expression of homeobox genes. In contrast, TALE and NF-Y co-occupy a subset of genomic elements at ZGA that is selectively associated with early-expressed genes involved in transcriptional regulation. Accordingly, disruption of NF-Y or TALE function produces phenotypes that share some features – particularly anterior deformations – but that also show unique defects, such as hindbrain abnormalities in TALE-deficient embryos. Notably, the co-occupied genomic elements possess enhancer activity when tested in a transgenesis assay in vivo. We conclude that TALE and NF-Y co-operate to control a subset of transcriptional regulators at ZGA, but also play distinct individual roles later in zygotic development.

## RESULTS

### NF-Y and TALE TFs are required for formation of anterior embryonic structures

In order to assess the roles of NF-Y and TALE TFs during embryogenesis, we first set out to disrupt their function. Previous work demonstrated that TALE TFs are required for formation of anterior embryonic structures, such that loss of various combinations of TALE factors results in animals with smaller heads, small (or absent) eyes, cardiac edema and hindbrain defects (Choe et al., 2002; Deflorian et al., 2004; Popperl et al., 1995; Popperl et al., 2000; Waskiewicz et al., 2001; Waskiewicz et al., 2002). This similarity in phenotypes is likely due to the fact that multiple TALE factors act together in larger protein complexes, which are rendered ineffective when one or more TALE factors are disrupted (reviewed in (Ladam and Sagerstrom, 2014; Merabet and Mann, 2016)). In preliminary experiments we recently observed abnormal anterior development also upon disruption of NF-Y function (Ladam et al., 2018). Here we extend this analysis to directly compare disruption of TALE factors (using the dominant negative PBCAB construct reported previously; (Choe et al., 2002)) to disruption of NF-Y function (using the previously reported NF-YA dominant negative construct; (Mantovani et al., 1994)) and find smaller heads in both cases (Figure 1A-D). A more detailed examination revealed abnormal head cartilage formation (53% of animals with disrupted NF-Y and 79% of animals with disrupted TALE function; Figure 1E-M) as well as loss of eyes (28% of animals with disrupted NF-Y and 19% of animals with disrupted TALE function; Figure 1X). Using in situ hybridization to detect expression of *pax2* (at the midbrain-hindbrain boundary), *krox20* (in rhombomeres 3 and 5 of the hindbrain) and *hoxd4* (in the spinal cord) in 24hpf embryos, we observed loss of r3 *krox20* expression upon TALE disruption (52% of embryos; as reported previously; (Choe et al., 2002; Deflorian et al., 2004; Popperl et al., 2000; Waskiewicz et al., 2001; Waskiewicz et al., 2002)), but did not detect any effects of NF-Y disruption (Figure N-U, Y). We conclude that both NF-Y and TALE function in formation of the anterior embryo and that TALE TFs have a distinct role in patterning of the hindbrain.

**Figure 1:**
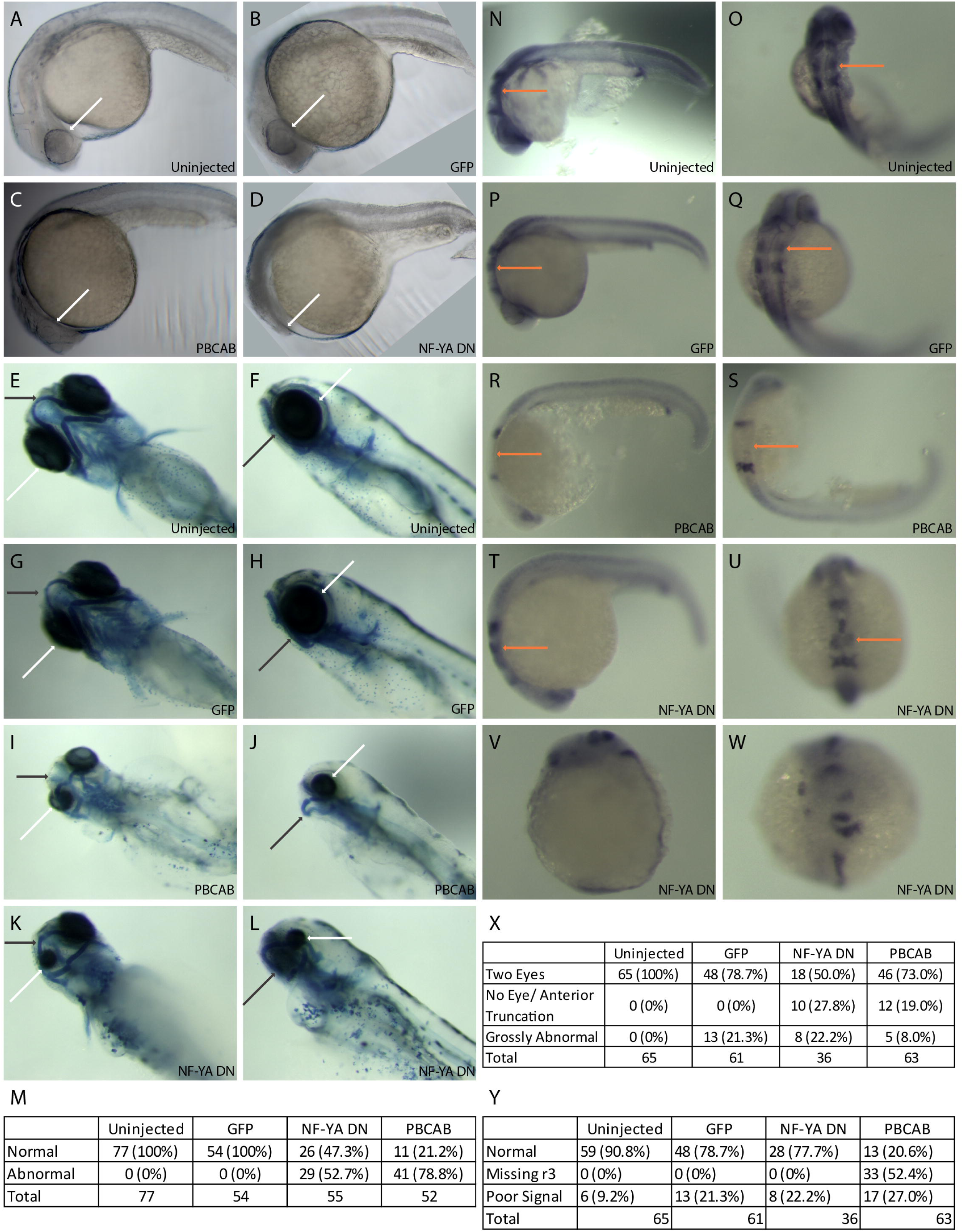
Disruption of NF-Y or TALE function affects anterior embryonic development. Zebrafish embryos were left uninjected (A, E, F, N, O) or injected with either control mRNA (GFP; B, G, H, P, Q), mRNA encoding a TALE dominant negative construct (PBCAB; C, I, J, R, S) or mRNA encoding an NF-Y dominant negative construct (NF-YA DN; D, K, L, T-W) at the 1-2 cell stage and raised to 24hpf (N-W), 28hpf (A-D) or 5dpf (E-L). Embryos were either left untreated (A-D), stained with alcian blue (E-L) or processed for detection of *pax2* (at the mid/hindbrain boundary), *krox20* (in rhombomeres 3 and 5) and *hoxd4* (in the spinal cord) transcripts by in situ hybridization (N-W). White arrows highlight differences in eye morphology (A-L), black arrows highlight differences in head cartilage formation (E-L) and orange arrows indicate differences in rhombomere 3 *krox20* expression (N-U). Tables summarize effects of TALE or NF-Y disruption on head cartilage formation (M), eye formation (X) and gene expression (Y). Panels V and W show representative images of embryos scored as having gross abnormalities in panel X. Embryos are shown in lateral (A-D, F, H, J, L, N, P, R, T, V) or dorsal (E, G, I, K, O, Q, S, U, W) views.

### NF-Y and TALE TFs have both shared and independent transcriptional targets

To identify shared and separate functions of the NF-Y and TALE TFs, we next carried out RNA-seq at 12hpf of zebrafish development (Figure 2A-C; Figure S1A-C). This timepoint was selected in order to ensure broad capture of gene expression changes resulting from disruption of NF-Y and TALE function. We find that disruption of TALE function affects the expression of 1,500 genes (646 downregulated, 854 up-regulated; FC≥1.5, p_adj_≤0.01; Figure 2B, D) at 12hpf. Since TALE factors are thought to act primarily as activators of transcription, we focused on genes downregulated upon disruption of TALE function. Applying the DAVID functional annotation tool, we find that TALE-dependent genes are enriched for functions related to transcription (particularly *hox* genes) as well as for factors controlling embryogenesis (Figure 2F; GO-terms for genes up-regulated upon disruption of TALE function are shown in Figure S1E). Accordingly, an examination of individual TALE-dependent genes identified members of several classes of TFs and developmental control genes (Figure 2G), consistent with the phenotype observed in figure 1. We next examined the effect of disrupting NF-Y function and find that 902 genes are affected (325 downregulated, 577 upregulated; Figure 2C, D) at 12hpf. An analysis of the GO terms associated with NF-Y-dependent genes revealed high enrichment in functions related to cilia and, to a lesser extent, in genes broadly controlling transcription and development (Figure 2H; GO-terms for up-regulated genes are shown in Figure S1F). In particular, different classes of TFs, as well as both structural and motor proteins found in cilia are downregulated upon disruption of NF-Y function (Figure 2I).

**Figure 2:**
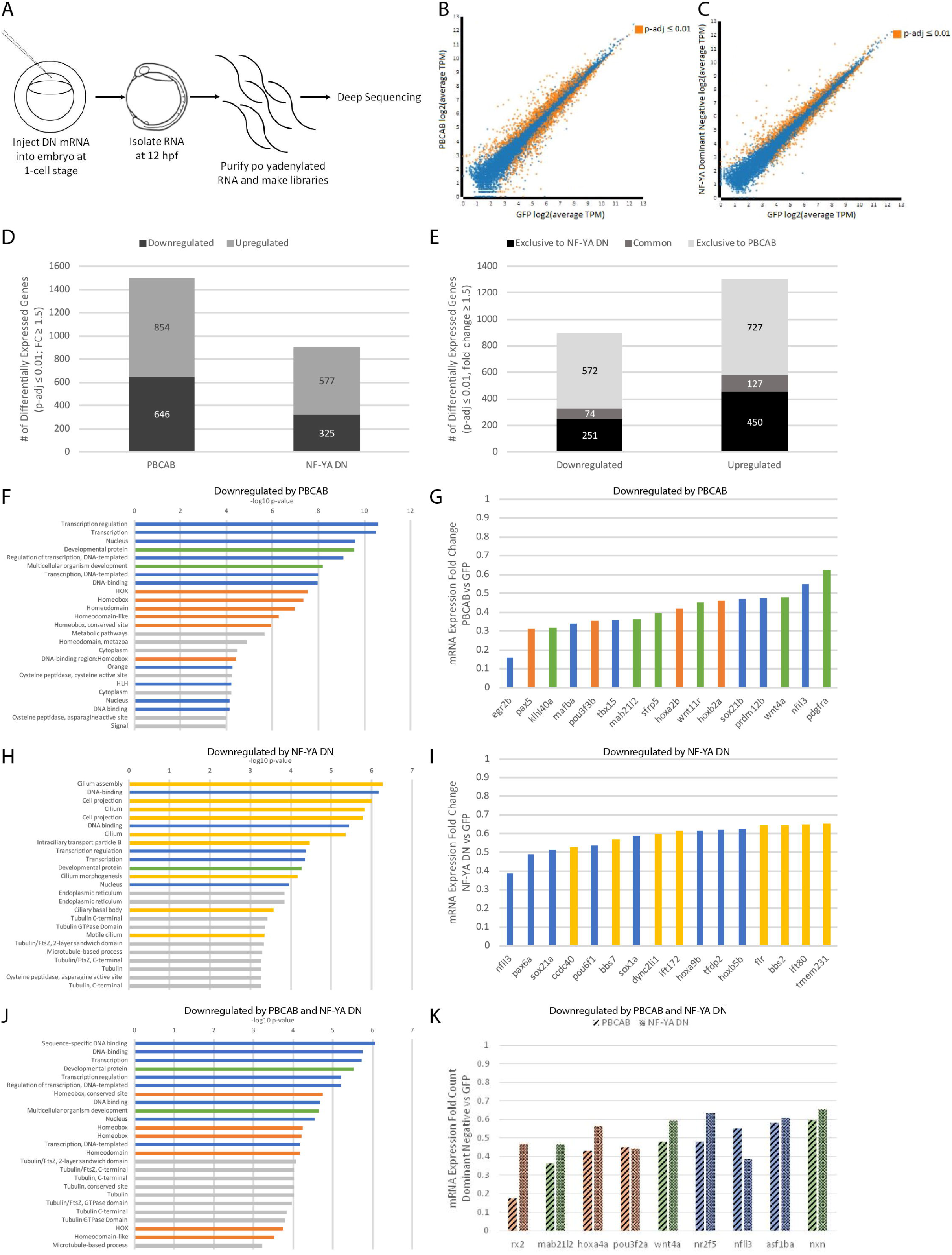
NF-Y and TALE TFs have both shared and independent transcriptional targets. (A) Schematic of RNA-seq experiments. (B-C) Scatterplots of gene expression in PBCAB vs GFP-injected (B) and NF-YA DN vs GFP-injected (C) zebrafish embryos (expression presented as log2 of average TPM for multiple replicates; see methods). Expression of genes highlighted in orange is significantly different at 12hpf (padj≤0.01; Wald test in DESeq2). (D) Number of genes differentially expressed in PBCAB (left) or NF-YA DN (right) relative to GFP-injected embryos (p-adj ≤ 0.01; fold-change ≥ 1.5). (E) Breakdown of downregulated (left) and upregulated (right) genes exclusive or common to each experimental condition. (F, H, J) DAVID analyses showing the 25 most significant GO terms (EASE Score) associated with genes downregulated by PBCAB (F), NF-YA DN (H), and common to both (J). Blue bars correspond to transcription-related, green to embryogenesis-related, orange to homeodomain-related, yellow to cilia-related, and gray to other ontologies. (G, I, K) Selected genes downregulated by PBCAB (G), NF-YA DN (I), or both (K). Color coding is the same as in (F, H, J). See also Figure S1.

Since disruption of either NF-Y or the TALE factors produces embryos with shared phenotypes, we next identified genes whose expression is dependent on both NF-Y and TALE function. We find that there are 201 such genes (74 downregulated, 127 upregulated; Figure 2E). Strikingly, the annotation of genes downregulated both upon disruption of TALE function and upon disruption of NF-Y function identifies transcriptional and developmental roles, but not roles associated with cilia (though several terms associated with tubulin function are retained; Figure 2J, K; GO-terms for up-regulated genes are shown in Figure S1G). We refer to this set of genes as ‘NF-Y/TALE co-regulated genes’ and note that it is a relatively small population, such that ∼23% (74/325) of the NF-Y dependent genes are also dependent on TALE function (Figure 2E). These results indicate that NF-Y and TALE TFs co-regulate a set of transcriptional and developmental control genes that is distinct from genes regulated by either NF-Y or TALE alone. Accordingly, a GO term analysis of genes regulated exclusively by NF-Y revealed strong enrichment for cilia genes (Figure S1I), while genes exclusively dependent on TALE function return GO terms enriched for transcriptional regulation such as *hox* genes (Figure S1H).

### NF-Y/TALE co-occupied genomic elements are distinct from those occupied by each TF separately

Given that NF-Y and TALE TFs appear to have both shared and independent functions, we next examined binding of these TFs across the zebrafish genome. In order to determine if these TFs have a role at ZGA, we focused our analysis at 3.5hpf – when zygotic genes are becoming active in zebrafish embryos. We previously used ChIP-seq to characterize Prep1 occupancy and found that this TF is bound at many genomic elements at maternally controlled stages (3.5hpf and earlier; (Choe et al., 2014; Ladam et al., 2018)), consistent with reports that TALE factors are maternally transmitted in zebrafish (Choe et al., 2002; Deflorian et al., 2004; Waskiewicz et al., 2002). Specifically, our analysis identified a 10bp motif (TGATTGACAG; termed the ‘DECA motif’; (Chang et al., 1997; Knoepfler and Kamps, 1997)) as the predominant element occupied by Prep1 in 3.5hpf zebrafish embryos. The DECA motif contains two half-sites – one for Pbx proteins (TGAT) and one for Prep factors (TGACAG) – and Pbx factors are known to form dimers with Prep proteins (reviewed in (Ladam and Sagerstrom, 2014)). Accordingly, using ChIP-qPCR we previously demonstrated that zebrafish Pbx4 occupied 11 of 12 tested DECA sites in 3.5hpf zebrafish embryos (Ladam et al., 2018). We have now extended this analysis to the entire zebrafish genome by performing ChIP-seq for Pbx4 at 3.5hpf (Figure 3A-C; Figure S2A). We find that the majority of Pbx4 peaks overlap with a Prep1 peak (94% overlap at FE≥10; Figure 3A, B, F; Figure S2B, C) and that the predominant sequence motif at Pbx4 binding sites is indistinguishable from the DECA motif observed at Pbx4/Prep1 co-occupied sites (Figure 3D, G). We also find that the distribution of Pbx4 peaks relative to TSSs is similar to that for Prep1 (Figure 3O), with ∼50% of all binding sites located within 30kb of an annotated promoter element (Nepal et al., 2013) in both cases, and that sites co-occupied by Pbx4/Prep1 show a similar distribution (Figure 3O). GO-term analyses (Figure 3E) revealed that genes associated with Pbx4 binding sites are enriched for the same transcriptional regulation and embryogenesis functions that we previously identified for genes associated with 3.5hpf Prep1 bound sites (Ladam et al., 2018). As expected, genes associated with Prep1/Pbx4 co-occupied sites return essentially the same GO terms (Figure 3H). Notably, a large number of Prep1 binding sites do not overlap with Pbx4 peaks (Figure 3F). While this could indicate that Prep1 has functions independent of Pbx4, it may also be a reflection of different affinities of the two antisera. Nevertheless, our observations indicate that Pbx4 binding takes place primarily at DECA sites in the context of Pbx4/Prep1 heterodimers at this stage of embryogenesis. Here we focused on the Prep1/Pbx4 co-occupied sites and refer to them as ‘TALE sites’.

**Figure 3:**
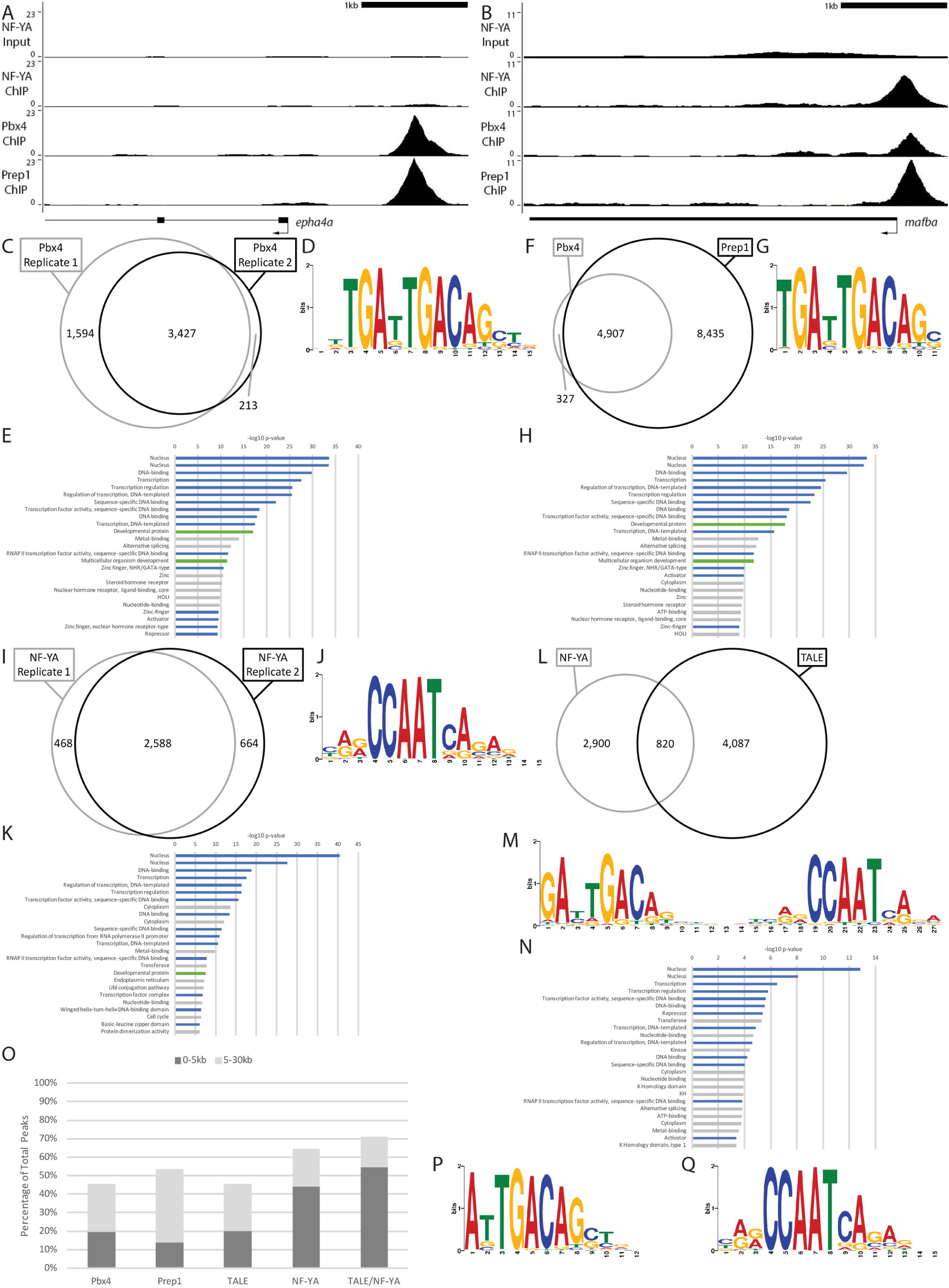
NF-Y/TALE co-occupied genomic elements are distinct from those occupied by each TF separately. (A-B) Representative UCSC Genome Browser tracks for NF-YA, Pbx4 and Prep1 ChIP-seq analyses at 3.5hpf. (C, F, I, L) Venn diagrams showing the overlap (at least 1bp shared between 200bp fragments centered on peaks) of two Pbx4 ChIP-seq replicates (C), the overlap of Pbx4 and Prep1 ChIP-seq peaks (F), the overlap of two NF-YA ChIP-seq replicates (I) and the overlap of TALE and NF-Y ChIP-seq peaks (L). (D, G, J, M) The top sequence motif returned by MEME for Pbx4-occupied sites (D), Pbx4/Prep1 co-occupied sites (G), NF-YA occupied sites (J) and TALE/NF-YA co-occupied sites (M). (E, H, K, N) The top 25 gene ontology (GO) terms returned by the GREAT analysis tool for genes associated with Pbx4-occupied sites (E), Pbx4/Prep1 co-occupied sites (H), NF-YA occupied sites (K) and TALE/NF-YA co-occupied sites (N). (O) Chart showing percent of ChIP-seq peaks found within 5kB or 30kB of a promoter. (P, Q) Top sequence motif returned by MEME for peaks bound by TALE, but not NF-YA (P) and peaks bound by NF-YA, but not TALE (Q). Only peaks with a 10-fold or greater enrichment over input (FE≥10) were considered for the analyses in C-I. See also Figure S2.

We previously reported that approximately 30% of TALE-occupied DECA sites observed at 3.5hpf have a CCAAT sequence motif nearby – usually at a distance of ∼20bp (Ladam et al., 2018). In other systems, such CCAAT boxes serve as binding sites for the heterotrimeric NF-Y transcription factor. Since NF-Y is maternally deposited in zebrafish (Chen et al., 2009), we previously used ChIP-qPCR to test 15 CCAAT boxes located near DECA sites and found that nine were occupied by NF-Y (Ladam et al., 2018). However, the commercial antiserum we used for the ChIP-qPCR experiment is too low affinity for ChIP-seq. Therefore, in order to examine NF-Y binding genome-wide, we raised antiserum to zebrafish NF-YA (the sequence-specific DNA binding component of the NF-Y heterotrimer; see Methods section) and carried out ChIP-seq for NF-Y on 3.5hpf zebrafish embryos (Figure 3A, B, I; Figure S2A). As expected, NF-Y-occupied genomic sites are highly enriched for the CCAAT box sequence motif (Figure 3J), but we find that the distribution of NF-Y peaks in the genome is somewhat different than the distribution of TALE peaks, such that NF-Y appears to be preferentially bound closer to promoters (Figure 3O). Further, we find that NF-Y-bound genomic elements are also associated with genes enriched for functions related to transcriptional regulation and embryogenesis (Figure 3K).

To further address the potential cooperation between NF-Y and TALE TFs, we next examined if NF-Y and TALE peaks co-localize in the zebrafish genome by determining if 200bp sequences centered on each peak overlapped by at least 1bp (see Methods section). Using this criterium, we find that approximately 22% of the NF-Y occupied sites overlap with a TALE-occupied site (corresponding to 17% of the TALE bound sites; Figure 3A, B, L; Figure S2B, C). Strikingly, motif analyses identified a ∼27bp sequence motif encompassing both a DECA motif and a CCAAT box (Figure 3M) associated with the NF-Y/TALE co-occupied sites, while sites occupied by TALE alone display a DECA motif (Figure 3P) and those occupied by NF-Y alone contain a CCAAT box (Figure 3Q). GO terms for genes associated with co-occupied sites are again enriched for functions related to transcriptional control, but less so for functions controlling embryogenesis (Figure 3N). Lastly, sites co-occupied by NF-Y and TALE factors show high association with promoters (Figure 3O). We conclude that, in the 3.5hpf zebrafish embryo, NF-Y and TALE TFs individually occupy genomic regions associated with both developmental and transcriptional regulators, but also co-occupy an extended binding motif that appears more selectively associated with transcriptional control genes.

### NF-Y and TALE co-regulate a subset of early-expressed transcriptional regulators

While our RNA-seq analysis identified genes regulated by NF-Y and TALE, it is not clear how direct this regulation might be. To begin addressing this question, we first examined whether TALE-dependent genes are associated with genomic elements bound by TALE TFs. We find that, of the 646 genes we identified as being TALE-dependent, 52% (335/646) are found near (as defined using default parameters in GREAT; see Methods section) a TALE occupied element (Figure 4A, B) and these genes are enriched for functions related to embryonic development and transcriptional regulation with a specific emphasis on *hox* genes (Figure 4C). Similarly, of the 325 genes our RNA-seq analysis showed to be downregulated upon disruption of NF-Y function, we find that 61% (199/325) are near a NF-Y occupied element (Figure 4A, B). The GO terms for these genes are enriched for functions related to transcriptional regulation, as well as for cilia structure/function (Figure 4D). Hence, 50-60% of NF-Y and TALE-dependent genes are associated with a binding site for the corresponding TF and the functional annotations of these genes show specific enrichment for the same terms as we observed in our RNA-seq analysis – cilia structure/function for NF-Y dependent genes and *hox* TFs for TALE-dependent genes. Since initial cilia formation and *hox* activity occurs at late gastrula and segmentation stages in zebrafish (Essner et al., 2002; Prince et al., 1998a; Prince et al., 1998b), these results indicate that NF-Y and TALE TFs control separate gene expression programs at these stages of zebrafish development.

**Figure 4:**
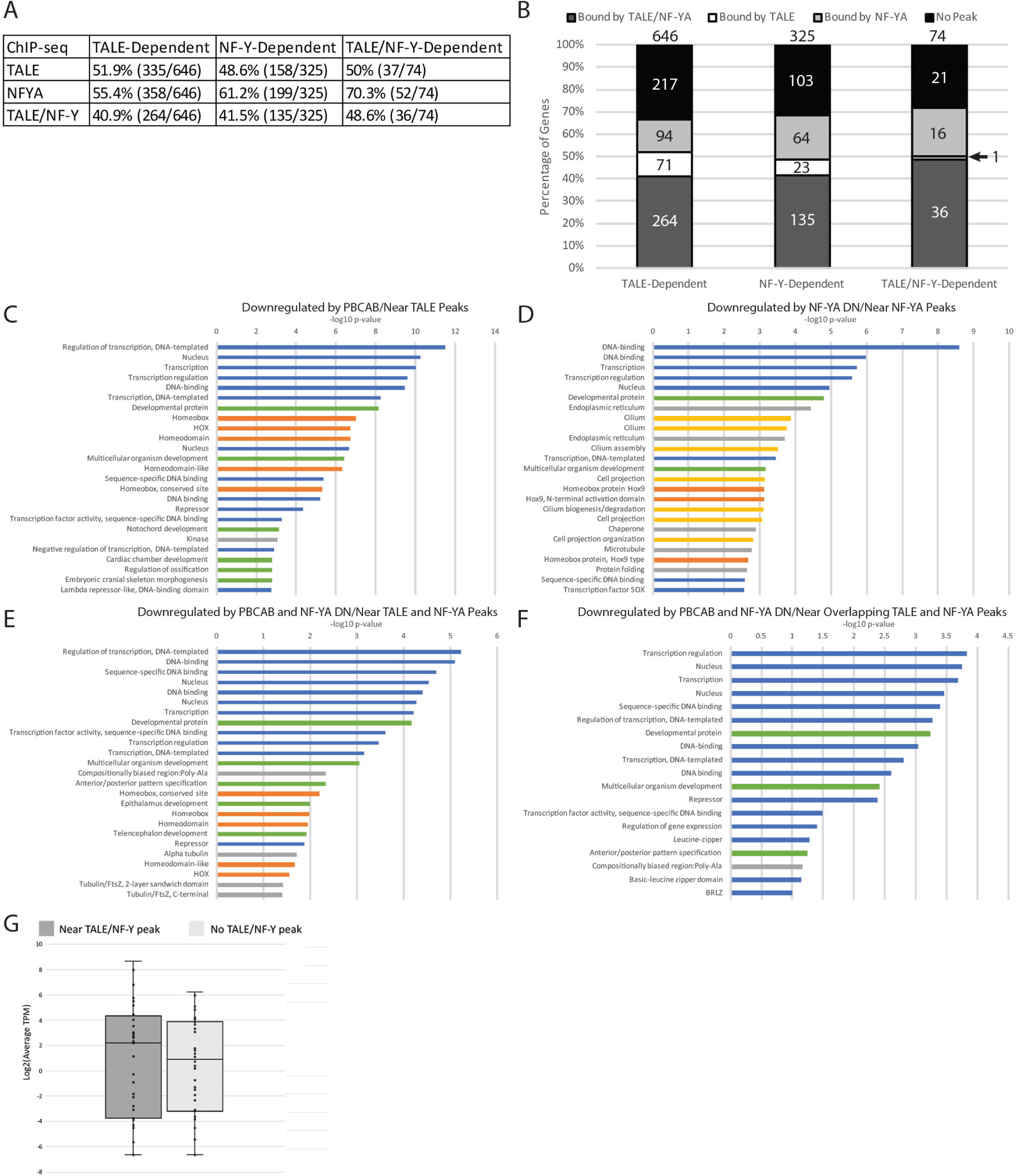
NF-Y and TALE co-regulate a subset of early-expressed transcriptional regulators. (A) Table showing correlation between NF-Y and/or TALE-dependent genes and binding by the corresponding TF at a nearby site. (ChIP peaks enriched by 4-fold or greater over input were considered). (B) Graphical breakdown of NF-Y and/or TALE occupancy near NF-Y and/or TALE-dependent genes. (C-F) Top GO terms returned by DAVID for TALE-dependent genes associated with TALE peaks (C), NF-Y dependent genes associated with NF-YA peaks (D), TALE/NF-Y dependent genes associated with both NF-YA and TALE peaks (E), and TALE/NF-Y dependent genes associated with overlapping TALE and NF-YA peaks (F). Blue bars correspond to transcription-related, green to embryogenesis-related, orange to homeodomain-related, yellow to cilia-related, and gray bars to other ontologies. (G) Box chart showing expression levels as log2(average TPM) for NF-Y/TALE-dependent genes with and without nearby TALE/NF-YA peaks.

In order to begin assessing co-regulation by NF-Y and TALE TFs, we carried out the reciprocal analysis. In doing so, we find that 55% of the 646 TALE-dependent genes are associated with an NF-Y occupied site (358/646) and 49% of the 325 NF-Y dependent genes are associated with a TALE-occupied element (158/325), indicating that TALE and NF-Y TFs co-regulate a subset of their target genes (Figure 4A, B). To examine this co-regulation further, we next focused specifically on the NF-Y/TALE co-regulated genes defined in figure 2E, J, K. We find that of the 74 genes in this category, 70% are associated with an NF-Y (52/74) occupied site and 50% (37/74) with a TALE-occupied site (Figure 4A, B). Indeed, 49% of co-regulated genes are found near both NF-Y and TALE occupied sites (36/74). Strikingly, the GO terms of co-regulated genes associated with both NF-Y and TALE-occupied elements converge on functions related to transcriptional regulation and embryonic development (Figure 4E). Further, if we specifically focus on genes that are near regulatory elements with overlapping NF-Y/TALE peaks (as defined in Figure 3L-N), we find that they function in transcription and regulation of development, but the categories related to cilia and homeobox functions are no longer represented (Figure 4F). Importantly, broad expression of transcriptional and developmental control genes is the first event to take place after ZGA. Accordingly, by analyzing RNA-seq data collected at 6hpf – shortly after ZGA – we find that genes associated with both NF-Y and TALE-occupied elements are expressed at higher levels than genes that lack such an association (Figure 4G). Hence, our results indicate that genes co-regulated by NF-Y and TALE act uniquely in transcriptional control of embryogenesis at ZGA, while NF-Y and TALE each controls a distinct gene expression program at later stages.

### Genomic elements co-occupied by NF-Y and TALE TFs act as enhancers in vivo

Although our data show that many genomic elements co-occupied by TALE and NF-Y are found near promoters, TALE TFs are known to act at enhancers (Ferretti et al., 2005; Ferretti et al., 2000; Grieder et al., 1997; Jacobs et al., 1999; Popperl et al., 1995; Ryoo and Mann, 1999; Tumpel et al., 2007). Further, while NF-Y was originally identified as acting at promoters (reviewed in (Maity and de Crombrugghe, 1998)), more recent work revealed an important role for NF-Y at tissue-specific enhancers (Oldfield et al., 2014). To explore these relationships in greater detail, we examined the chromatin state at NF-Y/TALE co-occupied elements and found that both H3K4me1 and H3K27ac (that mark enhancers and promoters) are highly enriched already at 4.3hpf and persist at 9hpf (Figure 5A-D). We note that elements bound by NF-Y alone and, to a lesser extent, TALE alone have the same characteristics. In agreement with NF-Y/TALE co-occupied elements driving gene expression at this stage of development, we also find a dramatic increase in H3K4me3 modifications (a mark of active promoters) between 4.3hpf and 9hpf (Figure 5E, F).

**Figure 5:**
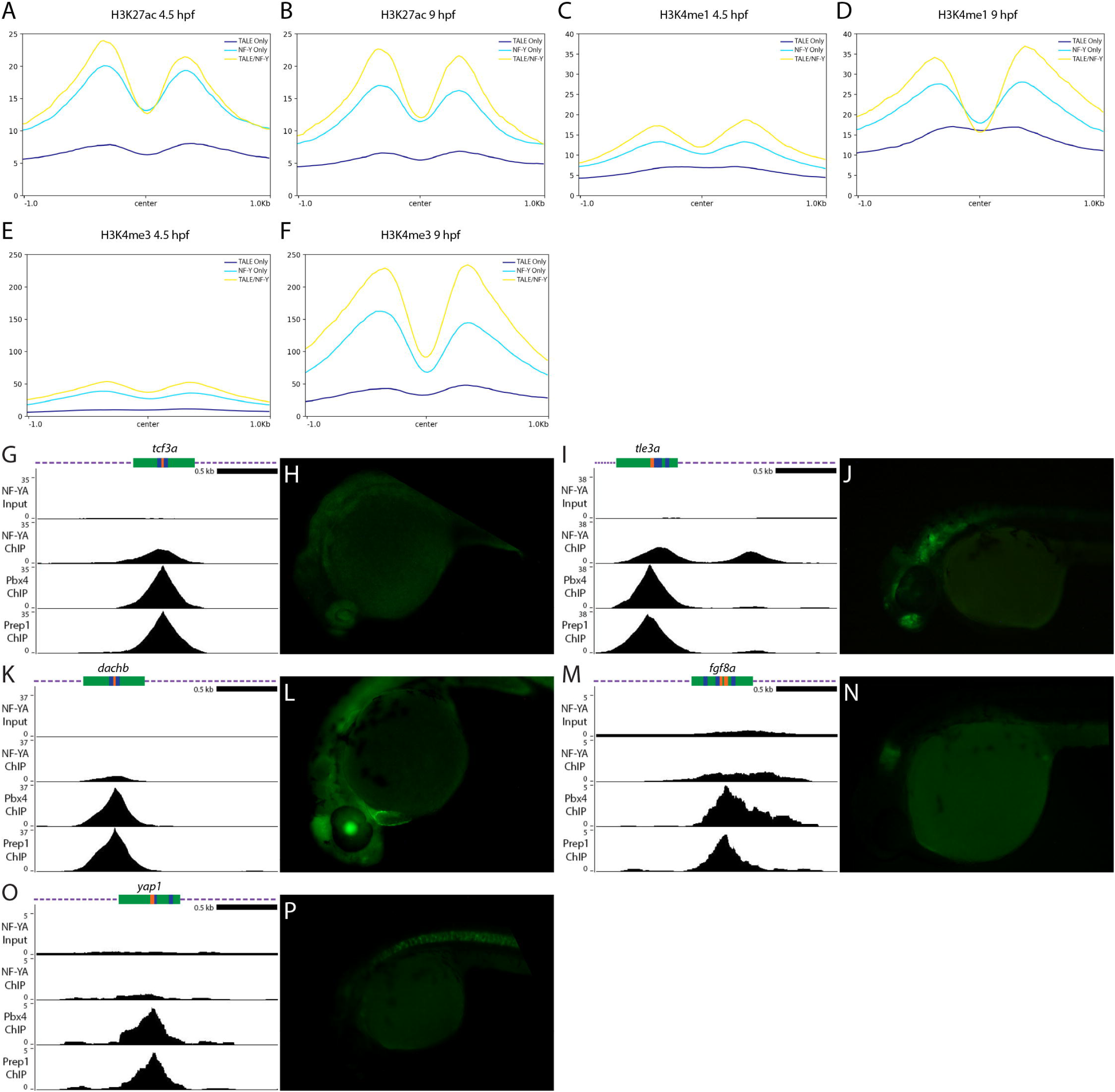
Genomic elements co-occupied by NF-Y and TALE TFs act as enhancers in vivo. (A-F) Average histone mark signals at genomic regions containing only TALE peaks (dark blue), only NF-YA peaks (light blue), or NF-YA/TALE peaks (yellow) for H3K27ac at 4.3hpf (A) and 9hpf (B), H3K4me1 at 4.3hpf (C) and 9hpf (D), H3K4me3 at 4.3hpf (E) and 9hpf (F). (G, I, K, M, O) UCSC Genome Browser tracks showing NF-YA, Pbx4, and Prep1 ChIP-seq data for the *tcf3a* (G), *tle3a* (I), *dachb* (K), *fgf8a* (M) and *yap1* (O) loci. The diagrams above the tracks show the putative enhancer region in green, DECA motifs in orange and CCAAT boxes in blue. (H, J, L, N, P) GFP expression in 24hpf F1 tcf3a:E1b-GFP (H), tle3a:E1b-GFP (J), dachb:E1b-GFP (L), fgf8a:E1b-GFP (N) and yap1:E1b-GFP (P) transgenic embryos resulting from crosses between male founders and wild type females. See also Figure S3.

To directly test if NF-Y/TALE co-occupied elements act as enhancers in vivo, we used a previously published enhancer assay (Li et al., 2010) and inserted individual genomic elements upstream of the E1b minimal promoter and the GFP reporter gene. We selected eight genomic elements that contain adjacent NF-Y/TALE motifs (as in figure 3N) and that are associated with genes expressed in the anterior embryo (Figure 5G, I, K, M, O; Figure S3N-Q) and used these to generate transgenic zebrafish. Of the eight constructs (named after the identity of the nearest gene), five showed expression in the F0 generation and GFP-positive embryos were raised to generate stable lines (Summarized in Figure S3Q). The remaining three constructs did not show F0 expression and were not considered further. In stable lines for each of the five constructs we detected tissue-restricted GFP expression with each construct producing a distinct pattern (Figure 5G-P). We screened at least two independent founders for each stable line and find that GFP expression is indistinguishable between founders carrying the same construct (Figure S3K, L, Q), indicating that each element imparts a unique tissue specificity to the basal E1b-GFP reporter that is independent of its integration site. In some instances, the observed expression pattern is comparable to that of the nearest gene (e.g. *fgf8a*; Figure 5N), suggesting that it represents an enhancer element controlling expression of the nearby gene. In other instances, the enhancer drives expression in a novel pattern (e.g. *yap1*; Figure 5P), suggesting that it may control a gene further away, or that the enhancer element tested (which is ∼500bp in length) lacks some inputs required for proper expression of the nearby gene. These results indicate that NF-Y/TALE co-occupied elements act as enhancers in vivo.

We next took two approaches to confirm that the observed expression patterns are dependent on NF-Y and TALE function. First, we expressed the dominant negative NF-Y and TALE constructs in embryos from a cross of the tcf3a:E1b-GFP transgenic line (Figure 6A). We find that GFP expression is dramatically reduced in embryos expressing either dominant negative construct (Figure 6B-E), indicating that expression from the *tcf3a* genomic element requires both TALE and NF-Y function. This observation was further confirmed by qRT-PCR analysis (Figure 6F). Second, we made use of a distinct transgenesis strategy that allows us to test the effect of mutating the TALE and NF-Y binding sites in a given enhancer element. Specifically, our transgenic construct includes the gamma-crystallin promoter driving GFP along with the candidate enhancer element driving RFP. We find that the wildtype *tcf3a* and *tle3a* enhancers drive tissue-specific RFP expression (Figure 6G, K), as expected based on our results in figure 5. However, when we test mutated versions of these elements (where the TALE and NF-Y binding sites have been disrupted), we find that transgenic animals (defined by GFP expression in the eye; Figure 6J, N) lack RFP expression (Figure 6I, M). We conclude that NF-Y/TALE co-occupied elements possess enhancer activity and that this activity requires NF-Y and TALE function.

**Figure 6:**
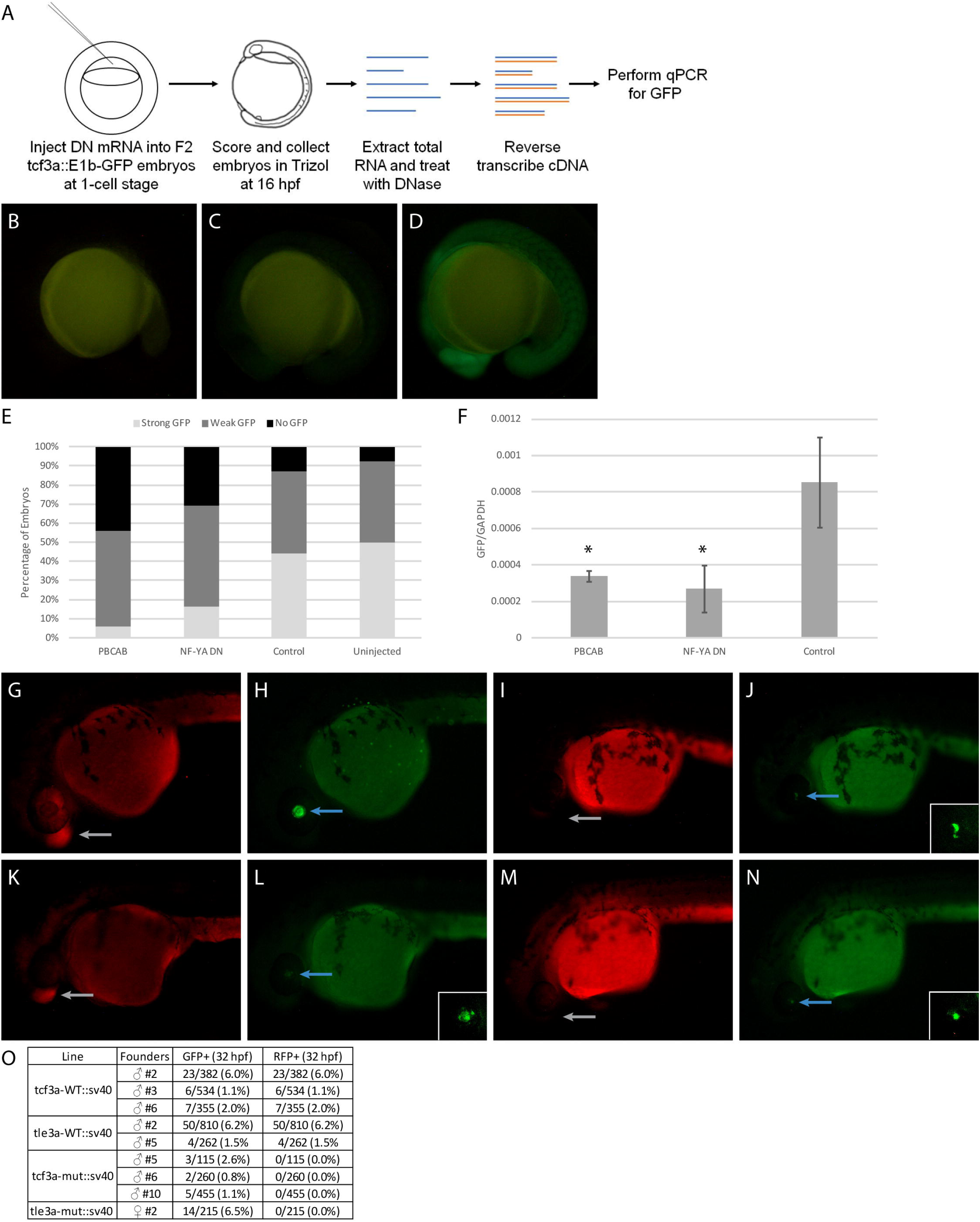
Disruption of TALE and NF-Y function reduces enhancer activity. (A) Schematic showing workflow for dominant negative disruption of tcf3a:E1b-GFP. (B-D) Representative images showing no GFP (B), weak GFP (C), and strong GFP (D) of dominant negative-injected embryos. (E) Distribution of GFP expression in uninjected embryos and embryos injected with PBCAB, NF-YA DN or control RNA. (F) RT-qPCR-based detection of GFP expression in embryos injected with PBCAB, NF-YA DN or control RNA. Data are shown as mean +/- SEM. Statistical test: unpaired t-test (G-N) Representative examples of RFP (G, K, I, M) and GFP (H, L, J, N) signal in tcf3a-WT:sv40 (G, H), tcf3a-mut:sv40 (I, J), tle3a-WT:sv40 (K, L) and tle3a-mut:sv40 (M, N) embryos at 32hpf. Insets in panels L, J, N show higher magnification of GFP expression in lens. Note that embryo in panels G, H is at a later stage than embryos in panels I-N. (O) Table quantifying results from experiment in panels G-N.

## DISCUSSION

Our results indicate that NF-Y and TALE TFs co-operate to regulate expression of transcriptional control genes active shortly after ZGA. These co-regulated genes are associated with an extended binding motif containing both TALE binding sites (DECA sites) and NF-Y binding sites (CCAAT boxes). While this motif has been observed previously (Penkov et al., 2013), it had not been assigned a function. This extended motif is also consistent with our previous finding that TALE and NF-Y proteins interact to form a complex (Ladam et al., 2018). Notably, the NF-Y and TALE TFs co-occupy these elements already at 3.5hpf – when ZGA is occurring. In fact, since NF-Y and TALE TFs are maternally deposited, they may be bound at these sites even earlier during embryogenesis. Accordingly, we have previously shown that Prep1 binding may be detected as early as 2hpf (Choe et al., 2014). Hence, co-occupancy by NF-Y and TALE marks a specific set of key early-acting genes during ZGA and likely even earlier during development. Given the apparently important function of NF-Y/TALE co-regulated genes, it is unclear why our disruption of NF-Y or TALE function does not produce a more severe phenotype. For instance, disruption of Nanog, SoxB1 and Oct4/Pou5f3, which are reported to act at ZGA in zebrafish, causes embryogenesis to stall at blastula stages (Lee et al., 2013). However, complete developmental blockade is achieved only when all three of these factors are disrupted, indicating that they act in combination to drive gene expression at ZGA. Additionally, recent work suggests that disruption of Nanog, which produces the most severe phenotype of the three TFs, may not affect ZGA directly, but instead block formation of essential extraembryonic tissues (Gagnon et al., 2018). Hence, it may be the case that vertebrate ZGA requires the action of multiple TFs and that disruption of any one TF is not sufficient to block ZGA. In agreement with such a model, recent work revealed that disruption of Dux, a TF implicated in murine ZGA, by itself does not block embryonic development (Chen and Zhang, 2019). Additionally, we find that NF-Y/TALE controls a relatively small number of genes at ZGA. This is in contrast to the situation in *Drosophila*, where the *Zelda* TF initiates broad gene expression at ZGA (Harrison et al., 2011; Nien et al., 2011), and suggests that multiple TFs may control subsets of genes at vertebrate ZGA.

Interestingly, we also find that NF-Y and TALE TFs act separately to regulate genes that have more specific functions at later stages of development. This is particularly clear for NF-Y, which regulates a cilia-related gene expression program, but also for the TALE TFs that appear preferentially associated with *hox* gene activity. Both of these processes are ongoing at late gastrula and organogenesis stages – accordingly, we detected gene expression changes in these programs by RNA-seq at 12hpf. While our ChIP-seq analyses indicate that genes in the cilia and *hox* programs are associated with NF-Y or TALE occupied elements – suggesting that they are directly regulated by NF-Y and TALE, respectively – we note that this ChIP-seq analysis was done at 3.5hpf. Hence, NF-Y and TALE TFs may remain bound at sites associated with genes in the cilia and *hox* programs, or new sites may be occupied at later stages. While we do not have ChIP-seq data at 12hpf for NF-Y, we have reported such data for Prep1 and find a large increase in the number of Prep1 binding sites from 3.5hpf to 12hpf. Importantly, these new sites are enriched near genes involved in *hox*-dependent functions, such as hindbrain development (Ladam et al., 2018). Hence, our data suggest that NF-Y and TALE co-operate at ZGA before taking on separate roles at later stages. We note that other putative ZGA regulators, such as Nanog and Sox TFs, also have later functions in embryogenesis, suggesting that this may be a generalizable concept.

Our results indicate that the genomic regions co-occupied by NF-Y and TALE TFs function as enhancers. As early as 4.3hpf (the earliest stage for which such data are available), these regions are enriched for histone modifications (H3K4me1 and H3K27ac) indicative of enhancer elements. In fact, in every case that one of the tested enhancers is transmitted by the mother, the resulting off-spring is GFP positive from the earliest stages of embryogenesis (Figure S3A-J) – suggesting that GFP is maternally deposited and that these enhancers may be active already in the maternal germline. H3K4me3 modifications (indicative of active transcription) are low at 4.3hpf, but increase by 9hpf, consistent with these enhancer elements being located near genes that are activated shortly after ZGA. Genomic elements occupied by either NF-Y or TALE also possess features indicative of enhancers, but, as discussed above, they are instead associated with later expressed genes. We also demonstrate that both NF-Y and TALE activity is required for the enhancer elements to drive gene expression in vivo, but it is not yet clear what specific functions are contributed by each TF. Previous work has suggested that both NF-Y and TALE may represent pioneer factors (Magnani et al., 2011; Nardini et al., 2013). Accordingly, we recently showed that many 3.5hpf TALE-bound sites are also occupied by nucleosomes (Ladam et al., 2018), suggesting that these TFs may be able to access their binding sites in nucleosomal DNA. Previous work has also demonstrated that TALE TFs can recruit histone modifying enzymes (Choe et al., 2014; Choe et al., 2009; Saleh et al., 2000) and may therefore promote the deposition of histone marks. Notably, we previously tested several of the NF-Y/TALE co-occupied elements for enhancer activity in HEK293 cells and failed to detect activity (Ladam et al., 2018). Also, both NF-Y and TALE are ubiquitously expressed during embryogenesis and therefore unlikely to mediate the tissue-specific expression we observe in the transgenic lines. Hence, it is possible that NF-Y and TALE are generally required for enhancer activity (possibly by rearranging nucleosomes and promoting histone modifications), but that additional tissue-specific TFs (that are present in embryos, but not in HEK293 cells) also act at these enhancers to drive expression in specific patterns.

## ACKNOWLEDGEMENTS

This work was supported by NIH grant NS038183 to CGS, BBSRC grant BB/N00907X/1 to NB and NIH grant DP3DK111898 to RM.

## AUTHOR CONTRIBUTIONS

Conceptualization, C.G.S., W.S., F.L. and N.B; Methodology, W.S. and C.G.S; Formal analysis, W.S., and I.J.D.; Investigation, W.S. and F.L.; Writing – Original Draft. C.G.S.; Writing – Review and Editing; W.S., F.L., I.J.D, T.J.P, R.M., N.B., and C.G.S.; Visualization, W.S.; Resources; T.J.P. and R.M.; Supervision, C.G.S., R.M., and N.B.; Project Administration, C.G.S.; Funding Acquisition, C.G.S., R.M., and N.B.

## STAR METHODS

### Lead Contact and Materials Availability

Further information and requests for resources and reagents should be directed to and will be fulfilled by the Lead Contact, Charles Sagerström.

RNA-seq data has been deposited in GEO under accession number GSE133459. ChIP-seq data has been deposited in ArrayExpress under accession number XX.

### Experimental Model and Subject Details

#### Care of zebrafish

The Institutional Animal Care and Use Committee (IACUC) of the University of Massachusetts Medical School approved all procedures involving zebrafish. Adult EkkWill zebrafish were maintained at 28°C in groups at a maximum density of 12 individuals per liter with constant flow. To collect embryos for timing-sensitive experiments, one adult male fish and one adult female fish were placed in separate chambers of a 500mL tank overnight then placed together the following morning for no more than 30 minutes. For experiments that were not timing-sensitive, both adults were placed in the same chamber overnight. Eggs were collected in 10cm dishes, immersed in egg water (60µg/mL Instant Ocean, 0.0002% methylene blue), and maintained in an incubator at 29°C. Dead and unfertilized eggs were manually removed after two hours.

### Method Details

#### Generation of mRNA for injection

Capped messenger RNAs encoding the dominant negative NF-YA (NF-YA DN; (Mantovani et al., 1994)), dominant negative Pbx4 (PBCAB; (Choe et al., 2002)) proteins were generated from 2µg of NotI-digested linearized pCS2+ plasmids using the mMessage mMachine SP6 Transcription Kit (ThermoFisher Scientific) according to the manufacturer’s guidelines. The RNA was purified using the RNeasy column with DNase treatment (Qiagen) according to the manufacturer’s guidelines. RNA quality was assessed on a 1% agarose gel and its concentration was measured on a NanoDrop instrument. 300pg of RNA injection mix containing water and 0.1% phenol red was injected into zebrafish embryos at the 1-cell stage. Injected embryos were raised to the proper stage according to animal care guidelines.

#### Characterization of TALE and NF-Y phenotypes

For gross phenotype assessment, 24hpf zebrafish embryos were placed on glass slides in 80% glycerol. For alcian blue staining, all incubations and washes took place on a nutator. 5dpf zebrafish embryos were fixed overnight in 4% phosphate-buffered paraformaldehyde. Following fixation, the embryos were washed in 0.1% phosphate-buffered Tween-20 (PBST) and bleached in 30% hydrogen peroxide for 2 hours. Once bleached, the embryos were rinsed twice in PBST and then stained overnight in alcian blue solution (1% hydrochloric acid (HCl), 70% ethanol, 0.1% alcian blue). After staining, the embryos were washed five times in acidic ethanol (HCl-EtOH; 5% HCl, 70% ethanol) with the final wash lasting 20 minutes. The embryos were then rehydrated in a series of 10-minute incubations of 75% HCl-EtOH/25% water, 50% HCl-EtOH/50% water, 25% HCl-EtOH/75% water, and 100% water and imaged. For in situ hybridizations, all incubations and washes took place on a nutator. 24hpf zebrafish embryos were fixed overnight in 4% phosphate-buffered paraformaldehyde. Following fixation, the embryos were washed in a 1:1 methanol:PBST solution, then PBST, and then treated with 1 µg/mL Proteinase K in PBST for 2 minutes. The embryos were washed once with −20°C acetone and twice with PBST then incubated at 70°C for 1 hour in Hyb+tRNA Buffer (50% formamide, 5X saline sodium citrate (SSC), 9.2mM citric acid, 0.5% Tween-20, 50 µg/mL heparin, 500 µg/mL tRNA). Next, the embryos were transferred to pax2/krox20/hoxd4a probe solution and incubated at 70°C overnight. After probe incubation, the embryos were washed sequentially for 10 minutes each at 70°C in Hyb Wash Buffer (50% formamide, 5X saline sodium citrate (SSC), 9.2mM citric acid, 0.5% Tween-20, 50 µg/mL heparin), 2:1 Hyb:2xSSC, 1:2 Hyb:2xSSC, 2xSSC, 0.2xSSC, and 0.1xSSC, then blocked in Blocking Solution (2% lamb serum and 2 µg/µL bovine serum albumin in PBST) at 4°C for 1 hour. The embryos were then incubated with 0.01% anti-DIG antibody at 4°C overnight. Following antibody treatment, the embryos were washed four times with PBST and two times with Staining Buffer (0.1M Tris pH 9.5, 50mM MgCl_2_, 125mM NaCl, 0.5% Tween20) then stained with Staining Solution (100 mg/mL polyvinyl alcohol, 0.35% 5-Bromo-4-chloro-3-indolyl phosphate, 0.45% 4-Nitro blue tetrazolium) at 37°C until the color developed. The embryos were then washed four times in PBST and scored. Sample size for phenotypic analyses was based on previous published reports that these dominant negative constructs produce phenotypes in >85% of injected embryos (Choe et al., 2002; Deflorian et al., 2004; Ladam et al., 2018; Waskiewicz et al., 2001). Embryos were randomly selected for inclusion in injected or control pools. No animals were excluded and experiments were not blinded.

#### RNA extraction

Zebrafish embryos were injected with either PBCAB, NF-YA DN, GFP, or antisense NF-YA DN mRNA as described above. At the desired timepoint, embryos were collected into three biological replicates of 50-100 embryos per condition. Dead animals were counted, but excluded from RNA extraction procedures. No other animals were excluded, and selection was not blinded. Each sample was placed in 1mL of Trizol and frozen at −80°C to help break up embryos. Once thawed, the embryos were broken up by pipette and 250µL of chloroform was added to each sample followed by vigorous shaking and a 3-minute incubation at room temperature. The samples were then centrifuged at 12,000*g for 15 minutes at 4°C and the aqueous phase was transferred to a new tube with 500mL of isopropanol and 10µg of GlycoBlue (ThermoFisher Scientific). The samples were vortexed, incubated at room temperature for 10 minutes, and then centrifuged at 12,000*g for 15 minutes at 4°C. The supernatant was removed, and the pellet washed in 75% ethanol then centrifuged at 7,500*g for 5 minutes at room temperature. The supernatant was once again removed, and the pellet was air-dried at room temperature for 10 minutes before resuspension in 50µL of water. The samples were then further purified and treated with DNase using the RNeasy Column kit (Qiagen) and eluted in 30µL of water.

#### RNA-seq library preparation and deep sequencing

The concentration and quality of each sample was assessed on a Bioanalyzer (Agilent), with all samples having a minimum RNA Quality Number of 8.0 and 28S/18S ratio of 1.0. 4µg of each sample of RNA was shipped to BGI for library preparation and sequencing. Polyadenylated RNAs were selected using oligo dT beads and then fragmented. N6 random primers were then used to reverse transcribe the library into double-stranded cDNA. A minimum of 20 million single-end 50bp reads were then generated with the BGISEQ-500 platform.

#### RT-qPCR

The concentration of each sample was assessed on a NanoDrop instrument. 1µg of RNA was reverse transcribed using the High Capacity cDNA Reverse Transcription Kit (ThermoFisher Scientific). To measure the quantity of select mRNAs, 25µL samples were prepared using 2µL of cDNA, 0.2mM of forward and reverse primer for each pair, and SYBR Green qPCR Master Mix (Bimake) according to the manufacturer’s guidelines. Measurements were made on a 7300 Real-Time PCR System (Applied Biosystems).

#### Generation of NF-YA Antiserum

Zebrafish NF-YA antiserum was prepared by ABClonal Technology. DNA encoding amino acids 1-328 of zebrafish NF-YA was cloned into the vector pET-28a-SUMO, containing a 12aa SUMO tag and a 6aa His tag. The vector was transformed into the *E. coli* Rosetta strain and the antigen peptide was induced with 0.8mM IPTG at 37°C for 4 hours. Small-scale antigen expression was confirmed by Western blot, showing a band at ∼58kDa corresponding to the peptide. The peptide was purified, appearing in both the supernatant and inclusion bodies. The concentration in the supernatant was 2mg/mL, which was deemed appropriate for immunization. Two rabbits were used for immunization and serum was collected on Day 52. The antiserum was tested by ELISA and deemed of sufficient quality with an OD450 > 0.4 at a 1:64,000 dilution. The antibody was purified via antigen affinity purification, with the polyclonal antibody concentration from animal #E7260 at 4.25mg/mL and from animal #E7621 at 4.66mg/mL. The antibodies were tested via Western blot at a 1:1000 dilution with 10, 5, 1, and 0.5ng of antigen. Bands of ∼60kDa were observed for antibodies from both animals at all four antigen concentrations.

#### ChIP-seq

Groups of ∼5,000 embryos (for Pbx4) and ∼10,000 embryos (for NF-YA) were collected at 3.5hpf and dechorionated in 1X pronase. The embryos were then dissociated by pipette, fixed in 2% formaldehyde in PBS for 10 minutes at room temperature, quenched with 125mM glycine, and flash-frozen in liquid nitrogen. Processing of cell pellets followed the protocol previously described (Amin et al., 2015). Nuclei were isolated in L1 Buffer (50mM Tris-HCl pH 8.0, 2mM EDTA, 0.1% NP-40, 10% glycerol, 1mM PMSF) then lysed in SDS Lysis Buffer (50mM Tris-HCl pH 8.0, 10mM EDTA, 1% SDS). Chromatin was sheared to an average length of 300bp using a Palmer immersion sonicator (Three 1-minute rounds of 10s on/2s off at 40% amplitude) and diluted 1:10 in ChIP Dilution Buffer (50 mM Tris-HCl pH8.0, 5 mM EDTA, 200 mM NaCl, 0.5% NP-40, 1 mM PMSF). The samples were pre-cleared with 50µL of Protein A Dynabeads (ThermoFisher Scientific) at 4°C for 3 hours, then Input samples were set aside and stored at −80°C. Next, 10µL of the appropriate antiserum was added (anti-Pbx4 or anti-NF-YA) and the samples were incubated rotating at 4°C overnight. The immune complexes were precipitated onto 50µL of Protein A Dynabeads, which were washed five times with Wash Buffer (20 mM Tris-HCl pH8.0, 2 mM EDTA, 500 mM NaCl, 1% NP-40, 0.1% SDS, 1 mM PMSF), three times with LiCl Buffer (20 mM Tris-HCl pH8.0, 2 mM EDTA, 500 mM LiCl, 1% NP-40, 0.1% SDS, 1 mM PMSF), and three times with TE Buffer (10 mM Tris-HCl pH8.0, 1 mM EDTA, 1 mM PMSF). To elute chromatin, the beads were incubated in 50µL of fresh Elution Buffer with shaking at 1,500 RPM for 15 minutes at 25°C then 15 minutes at 65°C. To reverse crosslinks, 2µL of 5M sodium chloride was added to the samples, which were then incubated at 65°C overnight. Purification of the DNA was accomplished using the MicroChIP Dia Pure Column kit (Diagenode) according to the manufacturer’s guidelines with an 11µL elution. To quantify the concentration of DNA, 1µL of each sample was passed through the dsDNA HS Assay (ThermoFisher Scientific) according to the manufacturer’s guidelines and quantified on a Qubit device.

#### ChIP-seq library preparation and deep sequencing

ChIP-seq libraries were prepared using the MicroPlex Library Preparation Kit v2 (Diagenode) according to the manufacturer’s guidelines. The entirety of each ChIP sample was used and Input samples were either diluted to the same concentration as their corresponding ChIP sample or, if the concentration of the corresponding ChIP sample was below the Qubit’s range, diluted to 0.2 ng/µL. Following library synthesis, an Illumina HiSeq4000 Sequencer was used to sequence the libraries.

#### E1b-GFP-Tol2 cloning

Putative enhancers of ∼500bp centered on Prep1 peaks near DECA sites and CCAAT boxes were amplified via PCR from 24hpf wild-type zebrafish genomic DNA using specific primers with XhoI sites (tcf3a, tle3a, dachb, fgf8a, pax5, her6, prdm14) or BglII sites (yap1) flanking either end (Table S1). The fragments were ligated into the E1b-GFP-tol2 (Birnbaum et al., 2012; Li et al., 2010) empty backbone digested with XhoI or BglII and transformed into competent DH5alpha E. coli cells (New England Biolabs). The amplified vector was validated by Sanger sequencing and purified using the Plasmid Maxi Kit (Qiagen).

#### Generation of pTransgenesis donor vectors

Mutant enhancers were generated by changing DECA sites contained within each enhancer to the sequence CGGTTGGTGC, which has been shown to prevent TALE binding (Vlachakis et al., 2000), and CCAAT boxes to the sequence ATGCG. Both mutant and wild-type versions of each enhancer were generated using gBlock technology (Integrated DNA Technologies). Due to limitations in gBlock synthesis, a 34bp AT-rich region at the 3’ end of the tcf3a enhancer could not be included compared to the E1b-GFP-tol2 version. A-tails were added to each end of the gBlock fragments using OneTaq Hot Start DNA Polymerase (NEB) (50ng of gBlock DNA, 1 unit of OneTaq Hot Start DNA Polymerase, 1X OneTaq Standard Reaction Buffer, 0.05mM dATP, 1.5mM MgCl_2_) and incubating the samples at 70°C for 30 minutes. 1µL of A-tailed gBlock fragment solution was then cloned into the pCR8 vector using the pCR8/GW/TOPO TA Cloning Kit (ThermoFisher Scientific) according to the manufacturer’s guidelines. The product was transformed into TOP10 chemically competent cells, validated by Sanger sequencing, and then purified using the Plasmid Midi Kit (Qiagen).

#### Generation of pTransgenesis vectors

pTransgenesis vectors were assembled using the LR Clonase II Plus enzyme mix (ThermoFisher Scientific). Four cassettes were assembled in one reaction, with gamma-crystallin:venusGFP as the p1 cassette (European Xenopus Resource Center (EXRC)), gBlock enhancers in pCR8 as the p2 cassette and Tol2/I-SceI-CH4-SAR/I-SceI/Tol2/P-element (EXRC) as the pDest-4 cassette. The p3.13 cassette was generated by ligating a BamHI/Bglll-digested gBlock (containing the SV40 minimal promoter) into BamHI-digested p3.13 Katushka-RFP plasmid (EXRC). 10fmol of each of the p1, p2, and p3 cassettes were combined with 20fmol of p4 cassette and 2µL of LR Clonase II Plus enzyme mix for the LR reaction in 10µL. The reaction was incubated at 25°C for 16 hours then treated with Proteinase K at 37°C for 10 minutes. 2µL of LR reaction product was transformed into Top10 chemically competent cells, validated by Sanger sequencing, and then purified using the Plasmid Maxi Kit (Qiagen).

#### Generation and observation of transgenic animals

Injection mixes containing 100ng/µL of E1b-GFP-Tol2 or pTransgenesis vector, 100ng/µL of Tol2 mRNA, and 0.1% phenol red were injected into wild-type zebrafish embryos at the 1-cell stage. The animals were observed for transient fluorescence for the first week, then raised to adulthood. Mature fish were crossed with wild-type fish and the offspring were observed for fluorescence. For E1b-GFP-Tol2 fish, GFP was observed as early as 18hpf. For pTransgenesis fish, RFP expression and GFP expression overlap was best observed at 32hpf, with RFP being apparent sooner and disappearing by ∼48hpf while GFP persisted after that time. Thus, any fish that appeared to be RFP+/GFP-were separated and observed for GFP expression at a later timepoint.

### Quantification and Statistical Analysis

#### RNA-seq analysis

RNA-seq analysis was performed using the University of Massachusetts Medical School Dolphin web interface. Ribosomal RNA reads were filtered out and FastQC was used to assess the quality of the remaining reads. RSEM_v1.2.28 with parameters -p4 --bowtie- e 70 --bowtie-chunkmbs 100 (Li and Dewey, 2011) was used to align the reads to the DanRer10 zebrafish transcriptome and normalize gene expression to transcripts per million (TPM). This revealed that PBCAB replicate 2 underperformed relative to the other samples and was excluded from further analysis. DeSeq2 (Anders and Huber, 2010) was used to identify differentially-expressed genes between three independent biological replicates of 12hpf embryos injected with GFP and three independent biological replicates of 12hpf embryos injected with NF-YA DN or between three independent biological replicates of 1 hpf embryos injected with GFP and two independent biological replicates of 12hpf embryos injected with PBCAB. DEBrowser was used to identify outliers among the replicates. To compensate for the exclusion of one replicate in GFP versus PBCAB analysis, only differentially expressed genes with a p-adj ≤0.01 (Benjamini and Hochberg FDR) were considered for analysis.

#### ChIP-seq Data Processing

All eight ChIP-seq fastq files (two independent 3.5hpf Pbx4 biological replicates, two independent 3.5hpf NF-YA biological replicates, and matched input DNA controls for each) contained 76bp paired-end sequences. The raw sequence quality was assessed with FastQC (https://www.bioinformatics.babraham.ac.uk/projects/fastqc/) and Fastq-screen (https://www.bioinformatics.babraham.ac.uk/projects/fastq_screen/). Next, remaining adapter reads were filtered out and poor-quality 3’ end sequences were trimmed with Trimmomatic version 0.36 (Bolger et al., 2014) using default parameters for ILLUMINACLIP and SLIDINGWINDOW and MINLENGTH set to 50bp. Using Bowtie2 version 2.2.3 (Langmead and Salzberg, 2012), the processed reads were then mapped to UCSC browser zebrafish genome release GRCz10 (danRer10/September 2014) (Tyner et al., 2017), and the mapped reads were further filtered with SAMtools view version 0.1.19 (Li et al., 2009) (with flags used -f 2 -q30) to remove reads with poor mapping quality and discordant mapped read pairs. To call peaks, the data, excluding reads that mapped to the mitochondrial genome and unassembled contigs in the assembly, was next passed through MACS2 version 2.1.0.20140616 (Zhang et al., 2008) with the q-value threshold set to 0.05 and default parameters except that the effective genome size was set to 1.03e9 (this equates to 75% of the total genome sequence, excluding ‘N’ bases).

#### ChIP-seq Analysis

Since the biological replicates for each factor demonstrated robust overlap, the sum of the two replicates was used for all subsequent analyses, by including all peaks meeting the selected cutoff in at least one of the biological replicates. Three different cutoffs were considered: all peaks with a fold enrichment (FE) ≥ 10, all peaks with a FE ≥ 4, and the top 10% of all peaks in each data set. The FE ≥ 10 cutoff showed the highest overlap between Pbx4 and Prep1 peaks as a percentage of the total Pbx4 peaks, and was selected as the best cutoff for ChIP-seq analysis (Supplementary Figure 3C). For a larger set of peaks, FE ≥ 4 peaks were considered for comparison to RNA-seq data (Figure 4).

#### ChIP peak overlap analysis

In the text, we use the term ‘overlap’ to indicate peaks identified as follows: ChIP peaks shared between different data sets were identified with the Intersect tool and exclusive peaks were identified using the Subtract tool in Galaxy (Goecks et al., 2010). All coordinates used were 200bp in length centered on peak summits and considered overlapping if they shared one or more base pairs.

#### qPCR Analysis

ddCt values were calculated from raw Ct values according to the formula 0.5^Ct^. Average ddCt values were then calculated by taking the mean of all three biological replicates. The ddCt of each GFP replicate was then normalized to the average gapdh ddCt according to the formula ddCt_GFP_/average ddCt_gapdh_ and then the mean of the normalized values was determined. Error bars were calculated based on the standard deviation of the three normalized GFP replicates in Excel. To determine whether the dominant negative conditions were significantly different from the control condition, an unpaired t-test was used in Excel, with p-values < 0.05 considered significant.

#### Determination of nearest genes to ChIP peaks

The number of Ensembl zebrafish transcription start sites within 5kb or 30kb of the summit of ChIP peaks was determined using the bedtools suite (Quinlan and Hall, 2010) in the Galaxy toolshed (Goecks et al., 2010). ChIP peak coordinates in danrer10 were converted to danrer7 (Zv9) using the LiftOver tool in the UCSC browser. The identities of genes near ChIP peaks were determined by the GREAT software version 3.0.0 (Hiller et al., 2013; McLean et al., 2010) using the he default settings of basal plus extension with proximal set to 5kb upstream and 1kb downstream and distal set to 1,000kb.

#### GO term analysis

Gene ontology (GO) terms enriched within different sets of genes were determined using DAVID version 6.8 (Huang da et al., 2009a, b). GO terms were ranked according to the EASE score, which was calculated based on a modified Fisher’s exact p-value, and graphed as the -log10 of that value.

#### DNA binding motif analysis

Significantly enriched binding motifs were identified using MEME and DREME within the MEME-Suite version 4.11.1 (Bailey et al., 2009; Machanick and Bailey, 2011). Both MEME and DREME were run according to their default settings. CENTRIMO was also run with default settings to determine the distribution of discovered motifs relative to ChIP peaks.

#### Chromatin feature analysis

Version 2.0 of the Deeptools (Ramirez et al., 2014) toolset in the Galaxy toolshed was used to create mean score profiles and heatmaps. Using the computeMatrix tool with region inputs of BED files containing ChIP coodinates and sample inputs of wiggle files from previously published data sets downloaded from GEO (Key Resources Table), signal matrices were generated in reference-point mode with the center set as the reference point. The distance upstream of the start sites and downstream of the end sites were set to 1000bp with a bin size of 25bp and ranked by mean signal when necessary. Heatmaps and profiles were generated from the matrices using the plotHeatmap and plotProfile tools respectively. The previously-published H3K27ac, H3K4me1, H3K4me3, and MNase data sets were all performed at 4.3hpf, which is somewhat later than the Pbx4, NF-YA, and Prep1 ChIP-seq experiments performed at 3.5hpf; however, asynchronous development in zebrafish embryos and large sample sizes make considerable overlap likely.

#### Data and Code Availability

RNA-seq data is available in GEO under accession number GSE133459 and ChIP-seq data is available in ArrayExpress under accession number E-MTAB-8137.

## Key Resources Table

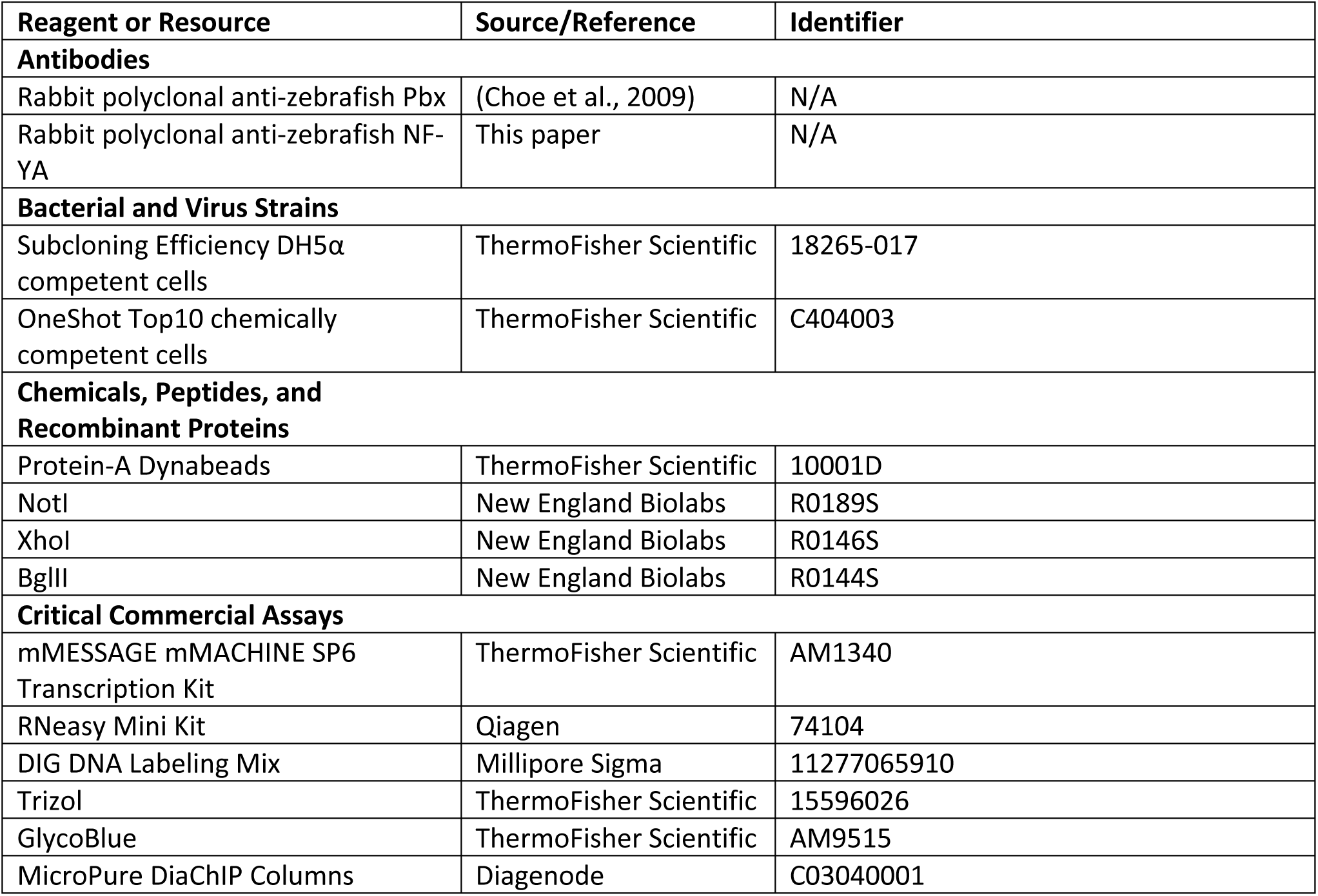
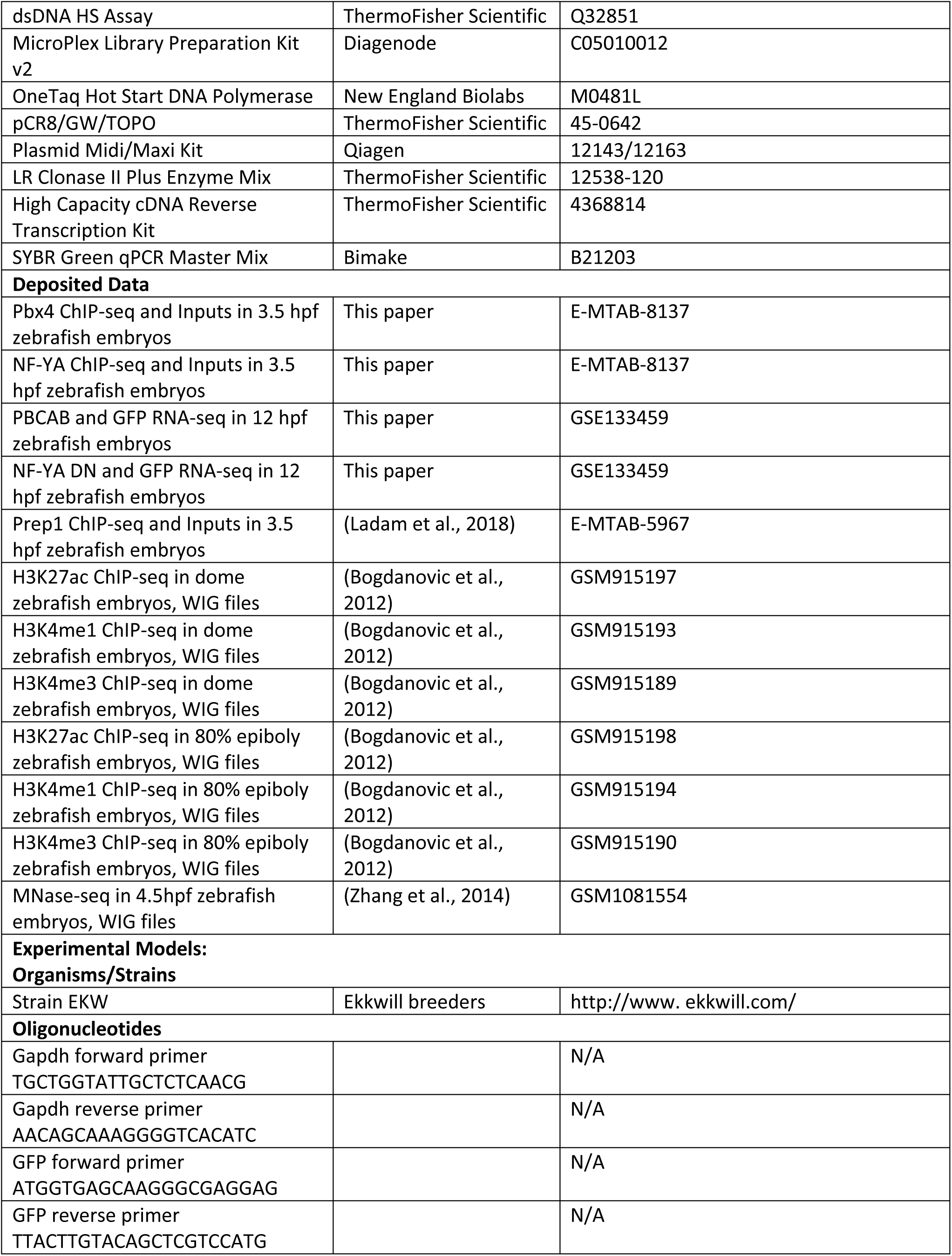
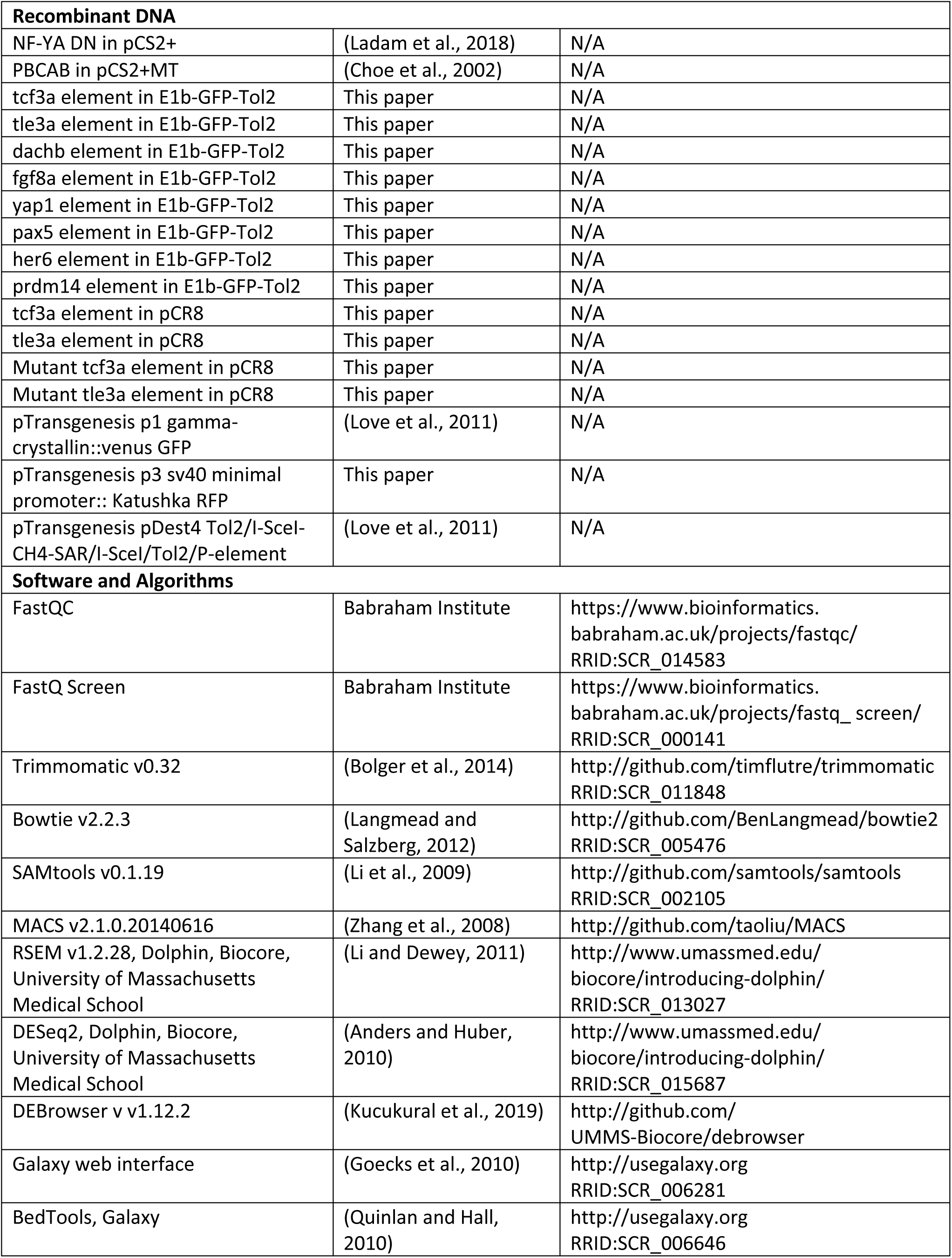
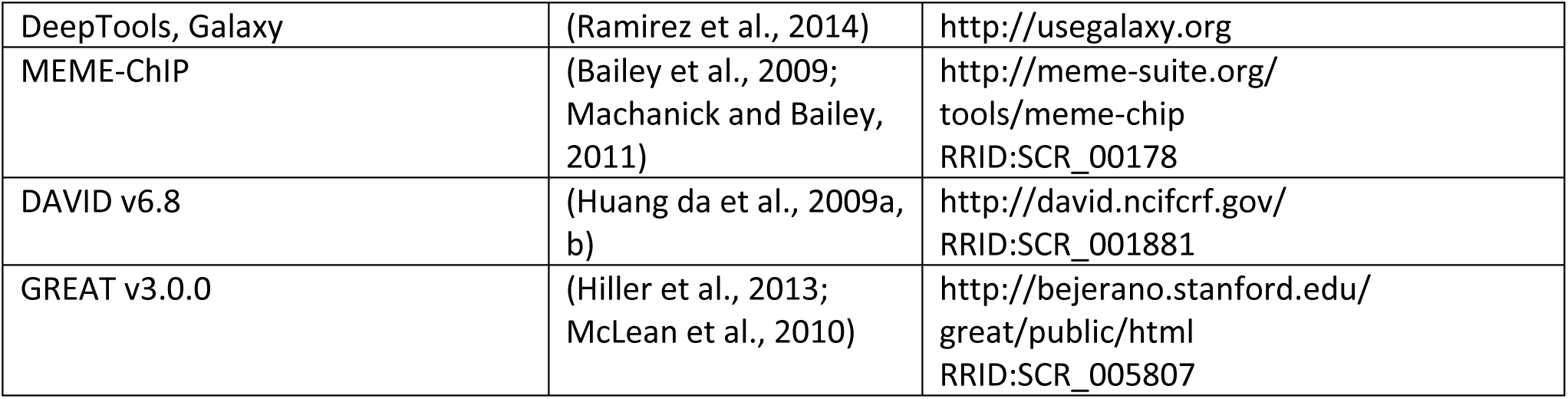

## SUPPLEMENTAL FILES

**Figure S1: Related to** Figure 2. **Identification of NF-Y and/or TALE-dependent genes in zebrafish.** (A) Read counts for the RNA-seq analysis. (B, C) Histograms, scatter plots, and Spearman’s rank correlation coefficient comparing each biological replicate of NF-YA DN with GFP (B) or PBCAB with GFP (C). (D) Venn diagram showing upregulated genes (p-adj ≤ 0.01; FC ≥ 1.5) in embryos injected with PBCAB or NF-YA DN. (E-I) GO terms associated with genes upregulated (p-adj ≤ 0.01, FC ≥ 1.5) by PBCAB (E), upregulated by NF-YA DN (F), upregulated by both PBCAB and NF-YA DN (G), downregulated exclusively by PBCAB (H) or downregulated exclusively by NF-YA DN (I). In E-I, blue bars correspond to transcription-related, green to embryogenesis-related, orange to homeodomain-related, yellow to cilia-related, and gray bars to other ontologies.

**Figure S2: Related to** **Figure 3. Identification of genomic binding sites for NF-Y, and Pbx4 in 3.5hpf zebrafish.** (A) Table showing data for Pbx4 and NF-YA ChIP-seq biological replicates with our previous Prep1 ChIP-seq data (Ladam et al., 2018) included as reference. (B) Table showing number of peaks that overlap (at least 1bp shared between 200bp fragments centered on peaks) between Prep1, Pbx4 and NF-YA ChIP-seq data sets. Only peaks with a 10-fold or greater enrichment over input were considered. (C) Table showing extent of overlap of Pbx4 peaks with Prep1 peaks and TALE peaks with NF-YA peaks at three different cutoffs (FE≥4, FE≥10 and top 10% of peaks).

**Figure S3: Related to** **Figure 5. Characterization of NF-Y/TALE-regulated enhancers in zebrafish.** (A-J) GFP-positive offspring from female transgenic carriers for *tcf3a:E1b-GFP* (A, B), *tle3a:E1b-GFP* (C, D), *dachb:E1b-GFP* (E, F), *fgf8a:E1b-GFP* (G, H), *yap1:E1b-GFP* (I, J) at 3.5hpf (A, C, E, G, I) and 24hpf (B, D, F, H, J). (K-L) GFP expression in F1 embryos from *yap1:E1b-GFP* founder #11 (K) and #5 (L). (M) Representative image of a 24hpf GFP-negative embryo. (N-P) UCSC Genome Browser tracks showing NF-YA, Pbx4, and Prep1 ChIP-seq data for the *pax5* (N), *her6* (O) and *prdm14* (P) loci. The diagrams above the tracks show the putative enhancer region in green, DECA motifs in orange and CCAAT boxes in blue. (Q) Table summarizing information about each putative enhancer element.

**Figure S4: Related to STAR methods section. Sequences of gBlocks used to generate pTransgenesis vectors containing putative enhancers.** Sequences are shown for gBlocks encoding wildtype and mutant tcf3a enhancers (A, B), wildtype and mutant tle3a enhancers (C, D) and the minimal SV40 promoter (E). In A-D, green highlights indicate wildtype DECA sites, blue indicate wildtype CCAAT boxes, purple indicate mutant DECA sites and red indicate mutant CCAAT boxes.

**Table S1: Related to STAR methods section. Primer sequences used to amplify putative enhancers from zebrafish genomic DNA.**

