## Supplemental Figures and Tables for "TALE and NF-Y co-occupancy marks enhancers of developmental control genes during zygotic genome activation in zebrafish"

### Figure S1

A

| Library | Total Reads | rRNA Reads | % rRNA | Reads - rRNA | Reads Aligned (RSEM) |
|---|---|---|---|---|---|
| NFYA DN1 | 25,570,095 | 118,566 | 0.46% | 25,451,529 | 15,618,541 |
| NFYA DN2 | 25,627,400 | 90,013 | 0.35% | 25,537,387 | 15,884,091 |
| NFYA DN3 | 25,648,288 | 103,952 | 0.41% | 25,544,336 | 15,847,491 |
| PBCAB1 | 22,951,454 | 6,211 | 0.03% | 22,945,243 | 14,084,935 |
| PBCAB2 | 25,649,514 | 7,074 | 0.03% | 25,642,440 | 15,668,814 |
| PBCAB3 | 25,660,266 | 7,991 | 0.03% | 25,652,275 | 15,605,294 |
| GFP1 | 25,565,467 | 4,756 | 0.02% | 25,560,711 | 16,034,166 |
| GFP2 | 25,488,663 | 4,725 | 0.02% | 25,483,938 | 15,765,669 |
| GFP3 | 25,696,572 | 6,392 | 0.02% | 25,690,180 | 15,980,155 |

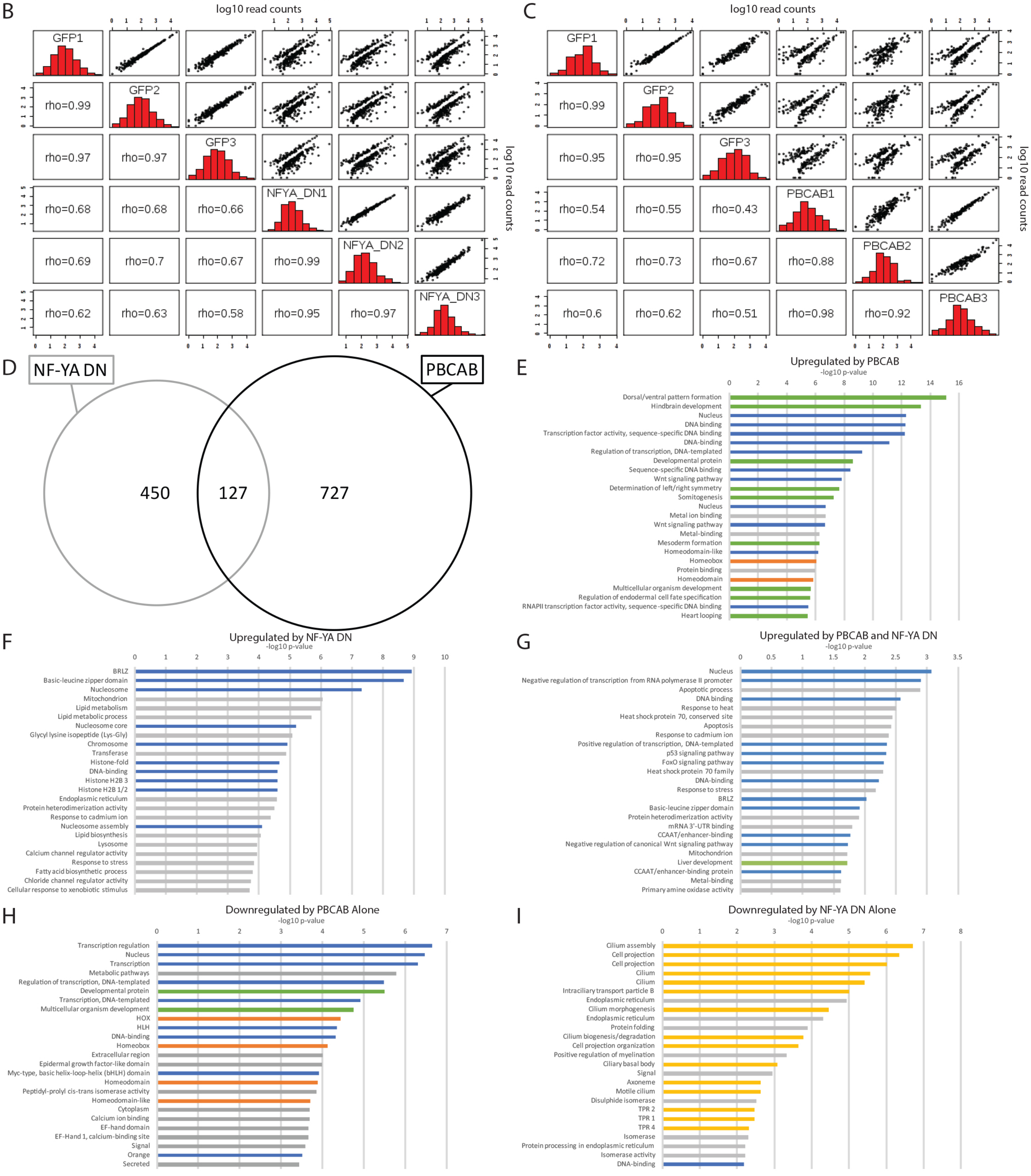

**Figure S1: Related to Figure 2. Identification of NF-Y and/or TALE-dependent genes in zebrafish.** (A) Read counts for the RNA-seq analysis. (B, C) Histograms, scatter plots, and Spearman's rank correlation coefficient comparing each biological replicate of NF-YA DN with GFP (B) or PBCAB with GFP (C). (D) Venn diagram showing upregulated genes (p-adj ≤ 0.01; FC ≥ 1.5) in embryos injected with PBCAB or NF-YA DN. (E-I) GO terms associated with genes upregulated (p-adj ≤ 0.01, FC ≥ 1.5) by PBCAB (E), upregulated by NF-YA DN (F), upregulated by both PBCAB and NF-YA DN (G), downregulated exclusively by PBCAB (H) or downregulated exclusively by NF-YA DN (I). In E-I, blue bars correspond to transcription-related, green to embryogenesis-related, orange to homeodomain-related, yellow to cilia-related, and gray bars to other ontologies.

### Figure S2

A

| Replicate | Total Peaks | FE≥10 Peaks | Common Peaks | FE≥10 Sum Peaks | Associated Genes (GREAT) |
|---|---|---|---|---|---|
| Pbx4-1 | 21,104 | 5,021 | 3,427 | 5,234 | 5,955 |
| Pbx4-2 | 34,119 | 3,640 | | | |
| Prep1-1 | 12,256 | 2,058 | 2,058 | 13,342 | 9,798 |
| Prep1-2 | 47,831 | 13,328 | | | |
| NF-YA1 | 21,784 | 3,056 | 2,588 | 3,720 | 4,398 |
| NF-YA2 | 22,196 | 3,252 | | | |

B

| ChIP-seq Data Sets | Overlapping Peaks | Associated Genes (GREAT) |
|---|---|---|
| Pbx4/Prep1 | 4,907 | 5,701 |
| Pbx4/NF-YA | 834 | 1,183 |
| Prep1/NF-YA | 937 | 1,332 |
| Pbx4/Prep1/NF-YA | 820 | 1,161 |

C

| ChIP-seq Overlap | FE≥4 | FE≥10 | Top 10% |
|---|---|---|---|
| Pbx4/Prep1 | 74.8% (13,836/18,508) | 93.8% (4,907/5,234) | 85.7% (2,960/3,455) |
| TALE/NF-YA | 13.2% (2,014/15,270) | 22.0% (820/3,720) | 23.5% (612/2,599) |

Figure S2: Related to Figure 3. Identification of genomic binding sites for NF-Y, and Pbx4 in 3.5hpf zebrafish. (A) Table showing data for Pbx4 and NF-YA ChIP-seq biological replicates with our previous Prep1 ChIP-seq data (Ladam et al., 2018) included as reference. (B) Table showing number of peaks that overlap (at least 1bp shared between 200bp fragments centered on peaks) between Prep1, Pbx4 and NF-YA ChIP-seq data sets. Only peaks with a 10-fold or greater enrichment over input were considered. (C) Table showing extent of overlap of Pbx4 peaks with Prep1 peaks and TALE peaks with NF-YA peaks at three different cutoffs (FE≥4, FE≥10 and top 10% of peaks).

### Figure S3

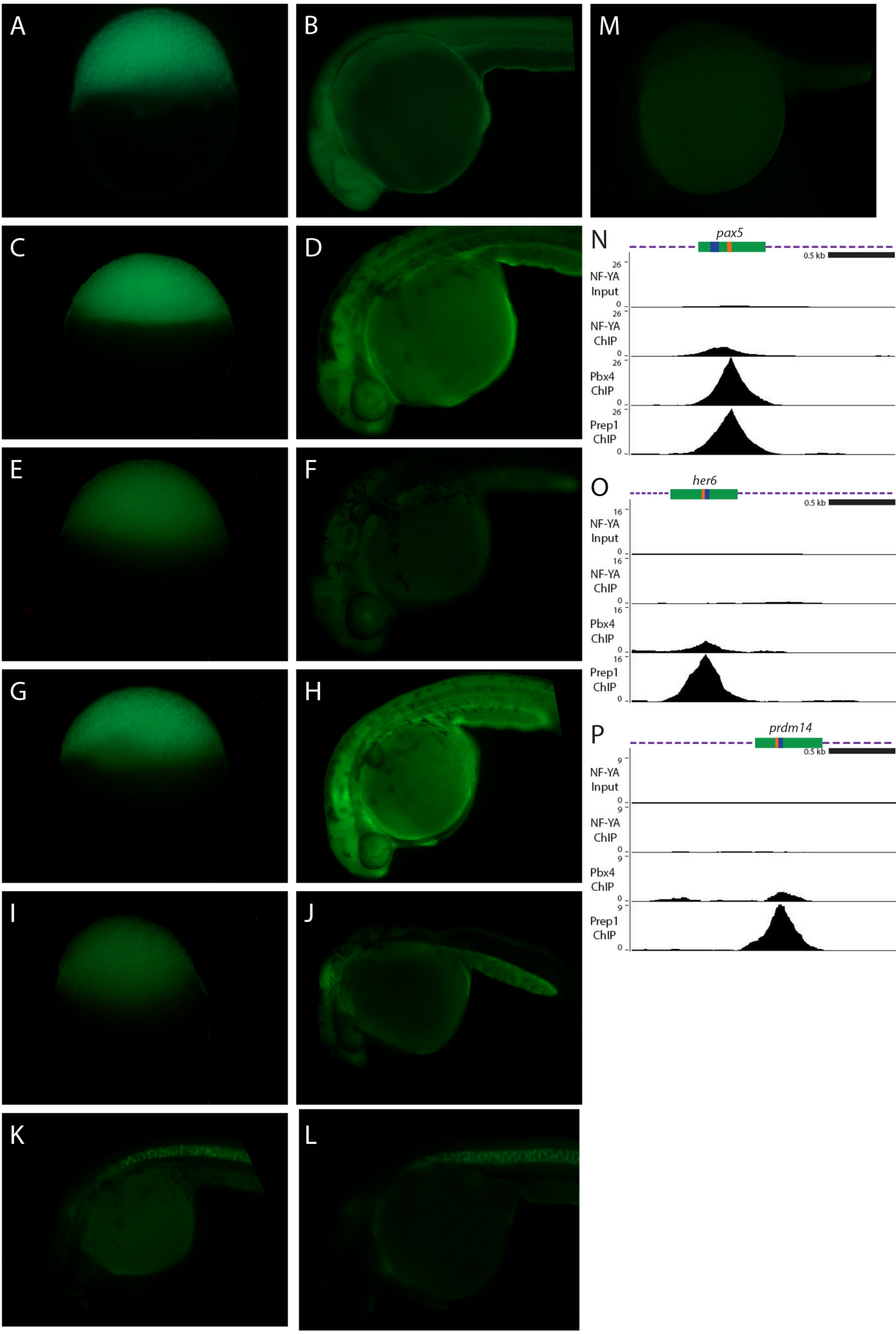


Q

| Element | Coordinates (GRCz10) | Size | NF-YA Peak | Pbx4 Peak | Prep1 Peak | Cloned in E1b::GFP | GFP+ Founders | GFP+ F1 0 hpf | GFP+ F1 24 hpf |
|---|---|---|---|---|---|---|---|---|---|
| tcf3a | chr2:57244797-57245296 | 500bp | Strong | Strong | Strong | Yes | ♀ #3 | 11/26 (42.3%) | 11/24 (45.8%) |
| | | | | | | | ♂ #4 | 0/84 (0.0%) | 20/84 (23.8%) |
| | | | | | | | ♂ #7 | 0/333 (0.0%) | 13/185 (7.0%) |
| tle3a | chr18:20235689-20236181 | 493 bp | Strong | Strong | Strong | Yes | | | |
| dachb | chr1:9669399-9669898 | 500 bp | Moderate | Strong | Strong | Yes | ♂ #6 | 0/364 (0.0%) | 26/308 (8.4%) |
| | | | | | | | ♂ #8 | 0/390 (0.0%) | 12/359 (3.3%) |
| | | | | | | | ♀ #15 | 25/198 (12.6%) | 25/174 (14.4%) |
| | | | | | | | ♀ #17 | 64/451 (14.2%) | 64/436 (14.7%) |
| fgf8a | chr13:28356309-28356808 | 500 bp | Moderate | Moderate | Moderate | Yes | ♀ #1 | 59/92 (64.1%) | 59/88 (67.0%) |
| | | | | | | | ♂ #10 | 0/77 (0.0%) | 31/72 (43.1%) |
| yap1 | chr18:37345724-37346223 | 500 bp | Weak | Moderate | Moderate | Yes | ♂ #4 | 0/221 (0.0%) | 33/156 (21.2%) |
| | | | | | | | ♂ #5 | 0/313 (0.0%) | 47/218 (21.6%) |
| | | | | | | | ♂ #11 | 0/25 (0.0%) | 3/21 (14.3%) |
| pax5 | chr1:21037015-21037514 | 500 bp | Moderate | Strong | Strong | Yes | 0 | | |
| her6 | chr6:36600313-36600812 | 500 bp | Weak | Moderate | Strong | Yes | 0 | | |
| prdm14 | chr24:14342429-14342928 | 500 bp | None | Weak | Moderate | Yes | 0 | | |

**Figure S3: Related to Figure 5. Characterization of NF-Y/TALE-regulated enhancers in zebrafish.** (A-J) GFP-positive offspring from female transgenic carriers for *tcf3a:E1b-GFP* (A, B), *tle3a:E1b-GFP* (C, D), *dachb:E1b-GFP* (E, F), *fgf8a:E1b-GFP* (G, H), *yap1:E1b-GFP* (I, J) at 3.5hpf (A, C, E, G, I) and 24hpf (B, D, F, H, J). (K-L) GFP expression in F1 embryos from *yap1:E1b-GFP* founder #11 (K) and #5 (L). (M) Representative image of a 24hpf GFP-negative embryo. (N-P) UCSC Genome Browser tracks showing NF-YA, Pbx4, and Prep1 ChIP-seq data for the *pax5* (N), *her6* (O) and *prdm14* (P) loci. The diagrams above the tracks show the putative enhancer region in green, DECA motifs in orange and CCAAT boxes in blue. (Q) Table summarizing information about each putative enhancer element.

**Figure S4:**

A: tcf3a-WT enhancer
TACTGCGTTAATCGCGCGTTTACTTTGATATTTAATCCACAACCAACACAATTAAAACGCCAAACATCAGC
GACGACAGTATATGTAACTTTATCCTGATATTTCCCGATTGTGCTTTAAATCACGCAGTACTAGACTCGCG
CGCGGAATGACACGACGCACTGTTGAAGAGCGATGGACTGAGAAAAAGTGCGAGATGGCACGATAGA
CCCACTGAGCGGACCAATAGCGATCGGGGAAAGTTTGATTGACGTATTCGGTGGCCAATCGAAGATCGT
GTTAACACGAAAGCCAAGCCTCTCTTCCATGCACACCCTAGCCAGGTTTTAAAAGAATGGCAACAGGAA
GCCATGGAATACTGTTGTGTTTTGTTGTTTGGTAAATGCTAATGTTTACCGCTAACCGCTCAAACTAACTT
CAAATGAATTCGACTCGAAACATAACATTGTTATTATTACATTTAGAC

B: tcf3a-mut enhancer
TACTGCGTTAATCGCGCGTTTACTTTGATATTTAATCCACAACCAACACAATTAAAACGCCAAACATCAGC
GACGACAGTATATGTAACTTTATCCTGATATTTCCCGATTGTGCTTTAAATCACGCAGTACTAGACTCGCG
CGCGGAATGACACGACGCACTGTTGAAGAGCGATGGACTGAGAAAAAGTGCGAGATGGCACGATAGA
CCCACTGAGCGGAATGCGAGCGATCGGGGAAAGTTCGGTTGGTGCATTCGGTGGATGCGCGAAGATCG
TGTTAACACGAAAGCCAAGCCTCTCTTCCATGCACACCCTAGCCAGGTTTTAAAAGAATGGCAACAGGA
AGCCATGGAATACTGTTGTGTTTTGTTGTTTGGTAAATGCTAATGTTTACCGCTAACCGCTCAAACTAACT
TCAAATGAATTCGACTCGAAACATAACATTGTTATTATTACATTTAGAC

C: tle3a-WT enhancer
ATAGATGACATTACCAGGACTGTATTGTTATATGGGTAACATGCGATTATGAGTGAGGGCTTTTTTTAAT
GTTATTAAGTGTTTGCATGCTCCTTTGCTCCTTTGTTTTATGTAAGGCTCTCATTACCACGTGGTAGTAAC
AGATTGTTTGAACTGGAAAGAAAAGCCATTCGAAGCTAATTAAGCAGCCATTCCAGGCACTATTCACGG
GCAGAAGAGCGAGAAGCACAGGCATTTGTCAGCGCTTGACCCCGCGTGGTATTGATTGACAACAAACCT
TCTTGAATGACAGCCTTAACCTTTCCCGTCCAATTGCAGTCGAGAGAATATAGATGCTGCTCTGCGATTG
GCTGAGAAGCTGTAAAGCCGCAAAGGGATCCCACGTGGGTGCAGCAGAAGAAACGGCACAGGATTGG
CCGCTTCTTCTGAGTTCAGACATGGCCGTTGTTCACGGAGATCAAACCTGAACAATCATCGTATTCCCAG
CGCTAGC

D: tle3a-mut enhancer
ATAGATGACATTACCAGGACTGTATTGTTATATGGGTAACATGCGATTATGAGTGAGGGCTTTTTTTAAT
GTTATTAAGTGTTTGCATGCTCCTTTGCTCCTTTGTTTTATGTAAGGCTCTCATTACCACGTGGTAGTAAC
AGATTGTTTGAACTGGAAAGAAAAGCCATTCGAAGCTAATTAAGCAGCCATTCCAGGCACTATTCACGG
GCAGAAGAGCGAGAAGCACAGGCATTTGTCAGCGCTTGACCCCGCGTGGTATTGATTGACAACAAACCT
TCTCGGTTGGTGCCCTTAACCTTTCCCGTATGCGTGCAGTCGAGAGAATATAGATGCTGCTCTGCGCGCA
TCTGAGAAGCTGTAAAGCCGCAAAGGGATCCCACGTGGGTGCAGCAGAAGAAACGGCACAGCGCATCC
CGCTTCTTCTGAGTTCAGACATGGCCGTTGTTCACGGAGATCAAACCTGAACAATCATCGTATTCCCAGC
GCTAGC

E: sv40 minimal promoter
aaagatctGCGATCTGCATCTCAATTAGTCAGCAACCATAGTCCCGCCCCTAACTCCGCCCATCCCGCCCCT
AACTCCGCCCAGTTCCGCCCATTCTCCGCCCCATCGCTGACTAATTTTTTTTATTTATGCAGAGGCCGAGG

CCGCCTCGGCCTCTGAGCTATTCCAGAAGTAGTGAGGAGGCTTTTTTGGAGGCCTAGGCTTTTGCAAAA
AGCTTGGCATTCCGGTACTGTTGGTAAAggatccaa

**Figure S4: Related to STAR methods section. Sequences of gBlocks used to generate pTransgenesis vectors containing putative enhancers.** Sequences are shown for gBlocks encoding wildtype and mutant tcf3a enhancers (A, B), wildtype and mutant tle3a enhancers (C, D) and the minimal SV40 promoter (E). In A-D, green highlights indicate wildtype DECA sites, blue indicate wildtype CCAAT boxes, purple indicate mutant DECA sites and red indicate mutant CCAAT boxes.

TABLE S1: Related to STAR methods section. Primer sequences used to amplify putative enhancers from zebrafish genomic DNA.

| **Primer** | **Sequence**[a] |
|---|---|
| Tcf3a-enh1F1 | ATGCCTCGAGTACTGCGTTAATCGCGCGTT |
| Tcf3a-enh1R1 | ATGCCTCGAGGTTAGTGTGATATAATCTGT |
| Tle3a-enh1F1 | ATGCCTCGAGGAAAAAAATAGATGACATTAC |
| Tle3a-enh1R1 | ATGCCTCGAGGCTAGCGCTGGGAATACGA |
| Dachb-enh1F1 | ATGCCTCGAGCGGTTTCTTTGCCATTCTTT |
| Dachb-enh1R1 | ATGCCTCGAGAACTAAGAACAATGTACG |
| Fgf8a-XhoIenhF1 | ATGCCTCGAGGGAGGTCGTTTGCGTATTTG |
| Fgf8a-XhoIenhR1 | ATGCCTCGAGCTTGTCAATCCACCCTGCTT |
| Yap1-enhF1 | |
| Yap1-enhR1 | |
| Pax5-enh1F1 | ATGCCTCGAGGCAAACGGATATTTTAAAAT |
| Pax5-enh1R1 | ATGCCTCGAGGTGCGTAAAAATCCAAGTAA |
| Her6-enh1F1 | ATGCCTCGAGTTCTTTTATAATTGTACTG |
| Her6-enh1R1 | ATGCCTCGAGTGATGTAAATAGAAATACTG |
| Prdm14-XhoIenhF1 | ATGCCTCGAGCCCCTCTTCTTTGTCCCTTG |
| Prdm14-XhoIenhR1 | ATGCCTCGAGGGTAGGCTATCTGGACGGATAAT |

[a] Each primer contains an XhoI site (CTCGAG) to allow it to be cloned into the E1b-GFP-Tol2 vector with the exception of yap1, which contained BglII sites (AGATCT).
